# ARPC5 Isoforms and Their Regulation by Calcium-Calmodulin-N-WASP Drive Distinct Arp2/3-dependent Actin Remodeling Events in CD4 T Cells

**DOI:** 10.1101/2022.01.25.477674

**Authors:** Lopamudra Sadhu, Nikolaos Tsopoulidis, Vibor Laketa, Michael Way, Oliver T. Fackler

**Affiliations:** Department of Infectious Diseases, Integrative Virology, University Hospital Heidelberg, Heidelberg, Germany; Department of Infectious Diseases, Virology, University Hospital Heidelberg, Germany; Center for Regenerative Medicine, Mass General Hospital, Harvard Medical School, Boston, MA, USA; Cellular Signalling and Cytoskeletal Function Laboratory, The Francis Crick Institute, London, UK; Department of Infectious Disease, Imperial College, London, UK

**Author notes:** Corresponding author at: Department of Infectious Diseases, Integrative Virology, University Hospital Heidelberg, Im Neuenheimer Feld 344, 69120 Heidelberg, Germany.

**Keywords:** Nuclear F-actin, Arp2/3 complex, CD4 T cell activation, DNA replication stress

## Abstract

Arp2/3-dependent formation of nuclear F-actin networks of different morphology and stability is observed in an increasing number of biological processes. In CD4 T cells, T cell receptor (TCR) signaling induces cytoplasmic and nuclear F-actin assembly via Arp2/3 to strengthen contacts to antigen presenting cells and to regulate gene expression, respectively. How Arp2/3 complex is regulated to mediate these distinct actin polymerization events in response to a common stimulus is unknown. Arp2/3-complex consists of 7 subunits where ARP3, ARPC1 and ARPC5 exist as two different isoforms in humans that can assemble in complexes with different properties. Examining whether specific Arp2/3 subunit isoforms govern distinct actin remodeling events in CD4 T cells, we find that the ARPC5L isoform drives nuclear actin polymerization, while cytoplasmic actin dynamics and TCR proximal signalling selectively relies on ARPC5. In contrast, formation of stable nuclear F-actin networks triggered by DNA replication stress in CD4 T cells requires ARPC5 and is independent of ARPC5L. Moreover, nuclear actin polymerization induced by TCR signaling but not by DNA replication stress is controlled by nuclear calcium-calmodulin signalling and N-WASP. Specific ARPC5 isoforms thus govern Arp2/3 complex activity in distinct actin polymerization events. ARPC5 isoform diversity thus emerges as a mechanism to tailor Arp2/3 activity to different physiological stimuli.

## Introduction

Development, proliferation and immune functions of T lymphocytes are regulated by their activation state. In concert with co-stimulatory receptors such as CD28, T-cell activation is primarily governed by engagement of surface exposed T-Cell Antigen Receptor (TCR/CD3) complexes with Major Histocompatibility Complex II-bound peptides on antigen-presenting cells (APC) (Valitutti, Dessing et al. 1995, Grakoui, Bromley et al. 1999, Sedwick, Morgan et al. 1999, Tskvitaria-Fuller, Rozelle et al. 2003, Dustin 2008). T cell activation triggers proliferation of already differentiated effector T cells but also drives polarized differentiation and proliferation of naïve T cells required for their development into effector T cells (Constant, Pfeiffer et al. 1995, Kaech, Wherry et al. 2002, Zhu and Paul 2008). Physiologically, APC-T cell interactions occur in the context of stable cell-cell contacts referred to as immunological synapse and trigger a broad range of downstream signaling events including sequential tyrosine phosphorylation cascades to activate PKC and MAPK signaling as well as rapid elevation of intracellular Ca^2+^ levels, which together regulates the expression of TCR target genes (Rhee and Choi 1992, Monks, Kupfer et al. 1997, Arendt, Albrecht et al. 2002, Dustin 2008, Oh-hora and Rao 2008, Joseph, Reicher et al. 2014, Monaco, Jahraus et al. 2016, Tsopoulidis, Kaw et al. 2019). TCR engagement also triggers the immediate polymerization of cortical actin at the immune synapse, which critically regulates downstream signaling by facilitating the formation and proper spatial distribution of signaling competent protein microcluster as well as by coordinating TCR and integrin signaling (Valitutti, Dessing et al. 1995, Morimoto, Kobayashi et al. 2000, Varma, Campi et al. 2006, Billadeau, Nolz et al. 2007, Jankowska, Williamson et al. 2018).

In addition to the cytoplasm, actin is also highly abundant in the cell nucleus, but the role of nuclear actin is much less studied. The advent of suitable probes to visualize nuclear F-actin facilitated the detection of nuclear actin assembly into complex structures (Baarlink, Wang et al. 2013, Melak, Plessner et al. 2017). Following the initial observation of short-lived F-actin networks in fibroblasts upon stimulation with serum, the rapidly growing field of nuclear actin polymerization has generated evidence for the formation of nuclear actin filaments in mammalian cells e.g. upon integrin and G-protein coupled receptor signaling or induction of DNA damage and during the repair of chromosome breaks, DNA replication stress, postmitotic nucleus expansion or upon infection with cytomegalovirus (Baarlink, Wang et al. 2013, Belin, Cimini et al. 2013, Wilkie, Lawler et al. 2016, Baarlink, Plessner et al. 2017, Caridi, D’Agostino et al. 2018, Schrank, Aparicio et al. 2018, Wang, Sherrard et al. 2019, Lamm, Read et al. 2020). The morphology of nuclear actin filaments ranges from stable thick cables to very transient thinner filament bundles. Different actin nucleators have been implicated in the formation of these meshwork: e.g. the formin Dia1 or Formin-1 / Spire1/2 for F-actin networks induced by serum and integrin-induced or DNA damage, or Arp2/3 complex for nuclear actin filaments induced in response to DNA damage (Baarlink, Wang et al. 2013, Belin, Lee et al. 2015, Caridi, D’Agostino et al. 2018, Wang, Sherrard et al. 2019, Lamm, Read et al. 2020).

Analyzing nuclear actin dynamics in response to TCR engagement in CD4 T cells, we previously observed the induction of a transient meshwork of thin nuclear actin filaments (Tsopoulidis, Kaw et al. 2019). This nuclear actin assembly, which drives a selective gene expression program required for the helper function of CD4 T cells, is triggered by nuclear calcium-calmodulin signaling and precedes actin polymerization in the cytoplasm. Interestingly, TCR-induced actin polymerization in the nucleus as well as in the cytoplasm relies on Arp2/3 complex but how spatio-temporal control of these distinct actin polymerization events is achieved is unclear. Arp2/3 is a multi-subunit complex that mediates actin polymerization in a wide range of diverse cellular processes, including the formation of lamellipodia, endocytosis and/or phagocytosis at the plasma membrane. Arp2/3 activity is regulated by interaction with nucleation promoting factors (NPFs) including WASP, N-WASP, WASH and WAVE (Rottner, Hänisch et al. 2010, Pizarro-Cerda, Chorev et al. 2017). NPF binding to and concomitant activation of Arp2/3 is induced by diverse upstream signals and the involvement of different NPFs therefore represents one level of spatio-temporal control of Arp2/3 complex. In addition, the three Arp2/3 subunits ARP3, ARPC1 and ARPC5 exist as two different isoforms in humans that can assemble in complexes with different biochemical properties (Jay, Berge-Lefranc et al. 2000, Millard, Behrendt et al. 2003, Abella, Galloni et al. 2016, von Loeffelholz, Purkiss et al. 2020, Galloni, Carra et al. 2021). As recently demonstrated for ARP3, different subunit isoforms can provide Arp2/3 complex with sensitivity for distinct upstream regulation (Galloni, Carra et al. 2021). To assess how Arp2/3 can mediate distinct actin polymerization events in the nucleus and cytoplasm in response to a common stimulus, we therefore investigated herein the role of Arp2/3 subunit isoforms and NPFs in these processes.

## Results

### Nuclear and plasma membrane actin polymerization triggered by CD4 T cell activation is sensitive to interference with Arp2/3 activity independently of the mode of activation

Nuclear and cytoplasmic actin remodeling in response to T cell activation can be visualized using several experimental approaches using Jurkat CD4 T cells stably expressing a nuclear lifeact-GFP reporter (JNLA) (Tsopoulidis, Kaw et al. 2019). Upon formation of an immune synapse between JNLA cells with Staphylococcus enterotoxin E (SEE) superantigen loaded Raji B cells (Fig 1A-D, Suppl. Video 1), actin polymerization is first observed in the nucleus and followed by actin polymerization at the cell-cell contact. These processes are recapitulated by surface-mediated stimulation of JNLA cells upon plating on dishes coated with anti-CD3 and anti-CD28 antibodies (Fig. 1E): TCR engagement rapidly induced a nuclear F-actin (NFA) meshwork, followed by cell spreading and actin polymerization into a circumferential F-actin ring (AR) at the cell periphery (Figs. 1F, G, Suppl. Video 3). The simultaneous visualization of nuclear and cytoplasmic actin dynamics is challenging due to their transient nature and occurrence in different focal planes. To specifically study nuclear actin polymerization, we therefore often induce T cell activation by PMA/Ionomycin (P/I), which only triggers nuclear actin polymerization in the absence of cytoplasmic actin remodeling or cell spreading (Tsopoulidis, Kaw et al. 2019) (Fig 1H-J). In all three experimental approaches, pretreating the cells with the Arp2/3 inhibitor CK869 prevents the formation of both NFA and, if induced by the stimulus used, AR with comparable efficacy (Figs.1C, D, G, J and Suppl. Video 2, 4). Arp2/3 complex thus mediates distinct phenotypically discernable actin polymerization events in CD4 T cell activation that are equally sensitive to pharmacological interference independently of the T cell stimulus used.

**Fig. 1.**
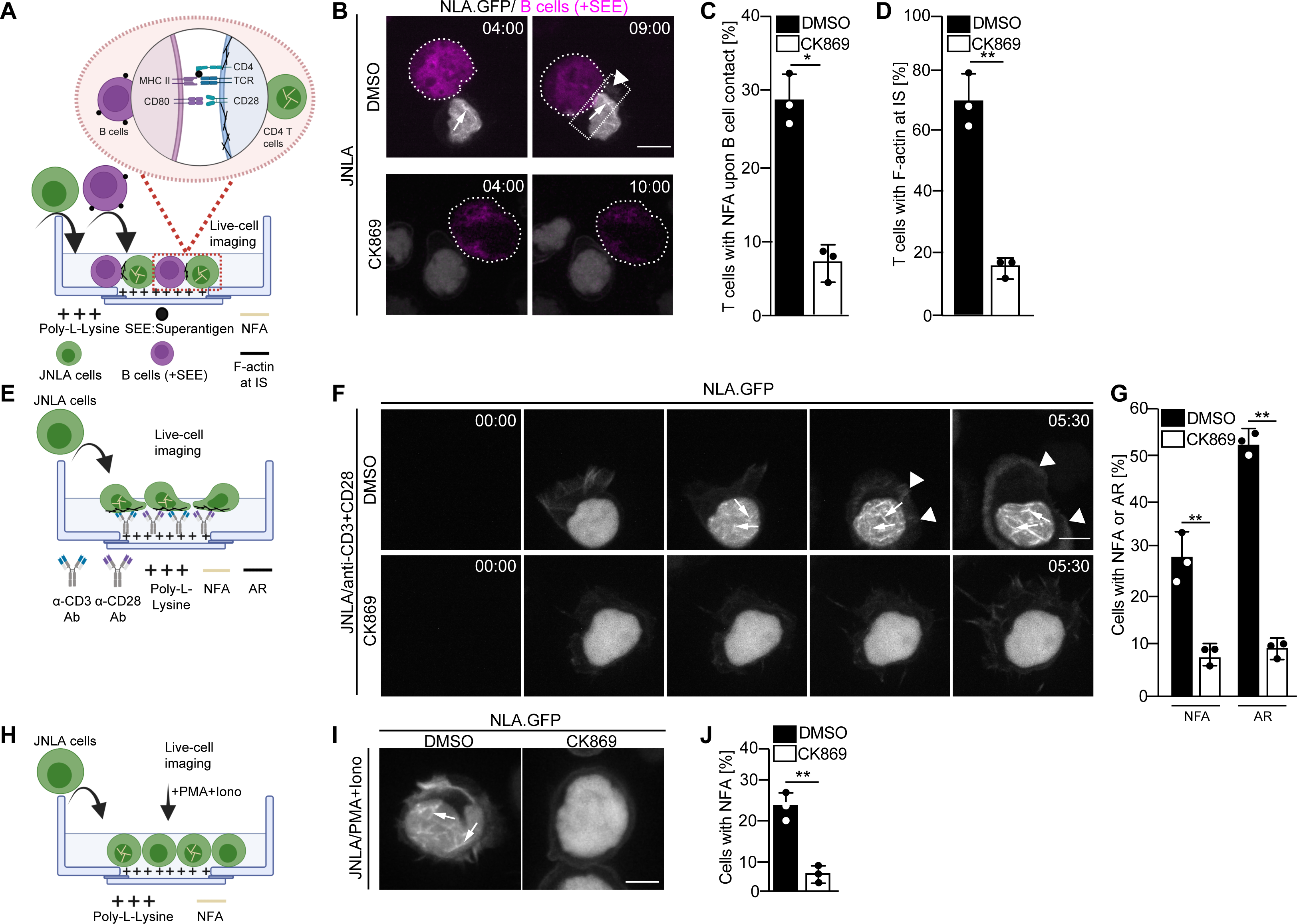
Arp2/3 complex mediated nuclear and plasma membrane actin polymerization in CD4 T cells. **(A)** Schematic representation of the experimental set up to visualize actin dynamics at the immune synapse, as performed for figures B-D. (**B**) Shown are representative still images at indicated time points from live-cell visualization of nuclear and plasma membrane actin dynamics in JNLA cells treated with either DMSO (solvent control) or CK869 upon contact with SEE pulsed Raji B cells. Images were acquired every 70s for a total of 30min after adding the Raji B cells. Still images represent the time at which the T and B cells made contact (left panel) to the time they formed an immune synapse (IS) as shown by the accumulation of plasma membrane F-actin at the contact site (right panel). Quantification of nuclear (**C**, NFA) and plasma membrane (**D**, AR) F-actin dynamics of JNLA cells upon contact with SEE pulsed Raji B cells are shown respectively. All data points indicate mean± s.d values from three independent experiments with at least 40 cells analyzed per condition per experiment. Scale bar, 7 µm. **(E)** Schematic representation of the experimental/live cell imaging set up on stimulatory GBDs as performed for F-G. (**F**) Jurkat CD4 T cells stably expressing nuclear lifeact-GFP (referred to as JNLA), pre-treated with either DMSO (solvent control) or CK869 for 30min, were put on TCR stimulatory GBDs and subjected to live-cell microscopy. Shown are representative still images from the spinning-disk confocal microscope from the time the cells fall on the coverslips until after contact with the stimulatory surface, with acquisition every 30s. Arrows indicate the nuclear F-actin (NFA) whereas arrowheads point to the F-actin at PM. Quantification of nuclear (**G:** nuclear actin filaments [NFA] and plasma membrane F-actin ring [AR]) polymerization are shown, respectively, upon contact with TCR stimulatory surface. Data points indicate mean±s.d values from three independent experiments where 40-60 cells were analyzed per condition in each experiment. **(H)** Schematic representation of the experimental/live cell imaging set up with P/I activation as performed for I-J. (**I**) JNLA cells pre-treated with either DMSO (solvent control) or CK869 for 30min, were put on poly-lysine coated GBDs and subjected to live-cell microscopy, which was then followed by addition of PMA+Ionomycin (P/I). Shown are representative still images from the spinning-disk confocal microscope forming NFA after P/I addition. Arrows indicate the NFA. (**J**) Quantification of nuclear actin polymerization upon addition of PMA+Ionomycin (P/I) was performed. Data points indicate mean values from three independent experiments where 40-60 cells were analyzed per condition in each experiment. Scale bar, 5µm. Nuclear f-actin (NFA) is denoted as yellow filaments within nucleus whereas plasma membrane f-actin is denoted as black filaments across all experimental schematics shown. Statistical significance based on the calculation of mean ± SD from three independent experiments, using Welch’s t-test were performed. \**P* ≤ 0.0332, \*\**P* ≤ 0.0021and *ns*: not significant.

### Expression of ARP2/3 isoforms in CD4 T cells

Nuclear but not cytoplasmic actin polymerization in response to T cell activation depends on nuclear Ca^2+^ transients and activation of nuclear Arp2/3 (Tsopoulidis, Kaw et al. 2019). In search for the molecular basis for this differential regulation of Arp2/3 complex, we hypothesized that nuclear and cytoplasmic actin polymerization involves distinct Arp2/3-complex subunit isoforms (Millard, Behrendt et al. 2003, Abella, Galloni et al. 2016). Analyzing the protein expression profile of Arp2/3 subunit isoforms in Jurkat CD4 T cells revealed expression of all Arp2/3 complex subunits (Fig. 2A), detection of ARPC5 however required extended exposure. Primary human resting CD4 T cells displayed low but detectable expression of all subunits and isoforms, with expression significantly increasing upon activation by anti-CD3/28 antibodies. This induction of protein levels by T cell activation was paralleled by induction of mRNA expression (Fig 2B: absolute values; Fig. 2C: relative to GAPDH; note that housekeeping genes are also subject to regulation by T cell activation (Sousa, Simi et al. 2019, Roy, McElhaney et al. 2020, Subbannayya, Haug et al. 2020). CD4 T cells thus express all ARP2/3 complex subunits isoforms and T cell activation increases Arp2/3 protein levels.

**Fig. 2.**
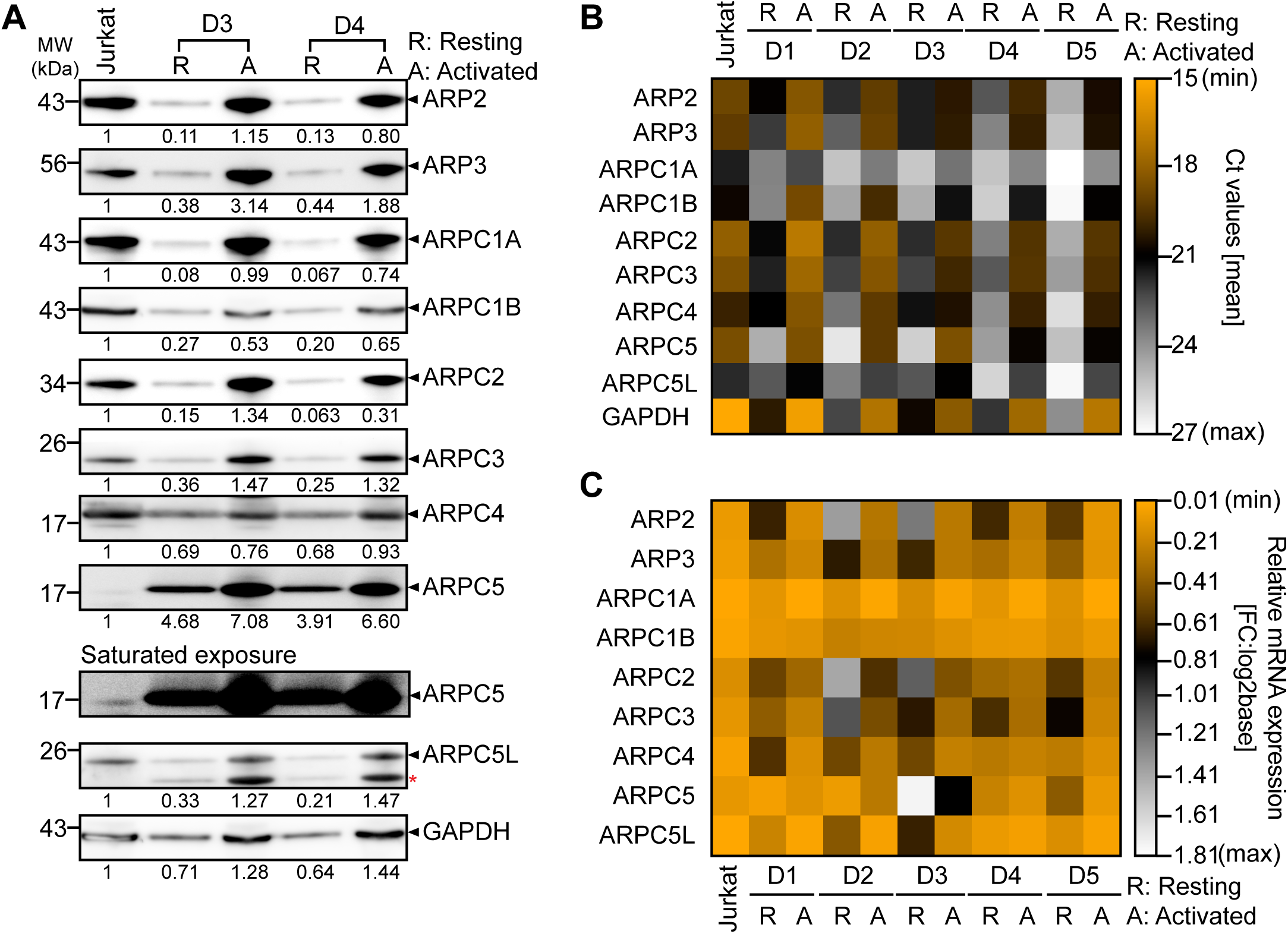
Expression of Arp2/3 complex-subunits and its isoforms. **(A)** Expression of all the subunits of the Arp2/3 complex along with the respective isoforms of ARPC1 and ARPC5 across Jurkat cell line and primary human CD4 T cells from two representative healthy donors were verified using Western blotting. Representative immunoblots compare the protein levels of each subunit and their isoforms in CD4 T cells. Additional comparisons for expression of these proteins in Resting (R) and Activated (A) CD4 T cells from Donor 3 and 4 are shown, respectively. Black arrowheads indicate the specific bands. Note that the ARPC5L antibody also detects ARPC5 (marked by red asterisk). The numbers indicated below each row represents mean±S.D values from three independent experiments of the densitometric quantification of the bands as compared to Jurkat protein levels (which is set to 1). Saturated exposure of the ARPC5 immunoblot is shown for better visualization of the ARPC5 levels in the JNLA cell line. **(B-C**) mRNA expression of all the subunits of the Arp2/3 complex along with the isoforms of ARPC1 and ARPC5 across Jurkat cell line and primary human CD4 T cells were additionally verified. The heatmap in **(B)** indicates the absolute differences in gene expression by comparing the Ct mean values obtained using qRT-PCR, for each subunit/isoform across Jurkat and primary human CD4 T cells. Colour code for the heatmap: yellow (high expression, low Ct mean values) and black-white (low expression, high Ct mean values). Whereas the heatmap in **(C)** shows the relative mRNA expression of each of the human Arp2/3 complex subunits and their isoforms, when normalized to the respective GAPDH levels of the cell line and primary CD4 levels and plotted as fold change (FC, as log of base2). Colour code for the heatmap: yellow (high expression, lowest difference after GAPDH normalization) and black-white (low expression, higher difference after GAPDH normalization). D1-D5 indicates CD4 T cells isolated from five different healthy human donors. Comparison of gene expression between RNA isolated from resting CD4 T cells (denoted here as Resting ‘R’) and from the CD4s that post activated for 72h with human anti-CD3+CD28 Dynabeads (denoted here as Activated ‘A’).

### Distinct ARPC5 isoforms mediate cytoplasmic and nuclear actin polymerization induced by TCR signaling

To assess the role of Arp2/3 subunit isoforms in the nuclear and cytoplasmic actin polymerization events triggered by TCR signaling, we transduced bulk JNLA cultures with isoform-specific shRNAs to reduce the expression of ARPC1A, ARPC1B, ARPC5 or ARPC5L (Fig. 3A). Selective silencing of ARPC1A, ARPC1B or ARPC5 did not significantly reduce the frequency of cells that displayed NFA in response to P/I. In contrast, JNLA cells with reduced ARPC5L levels were significantly impaired in NFA formation upon stimulation with P/I but also in response to TCR engagement (Figs. 3B, C, S1A, B). As with NFA, ARPC1A or ARPC1B were both able to support cell spreading and formation of cytoplasmic AR after surface mediated TCR stimulation (Figs. 3D-E, see Fig. S1C for a lower magnification overview and Figs. S1D-F for quantification of cell morphologies). Importantly, ARPC5 but not ARPC5L was required for efficient cell spreading and cytoplasmic AR assembly triggered by TCR engagement and cells lacking ARPC5 often displayed aberrant F-actin organization (Figs. S1C-D).

**Fig. 3.**
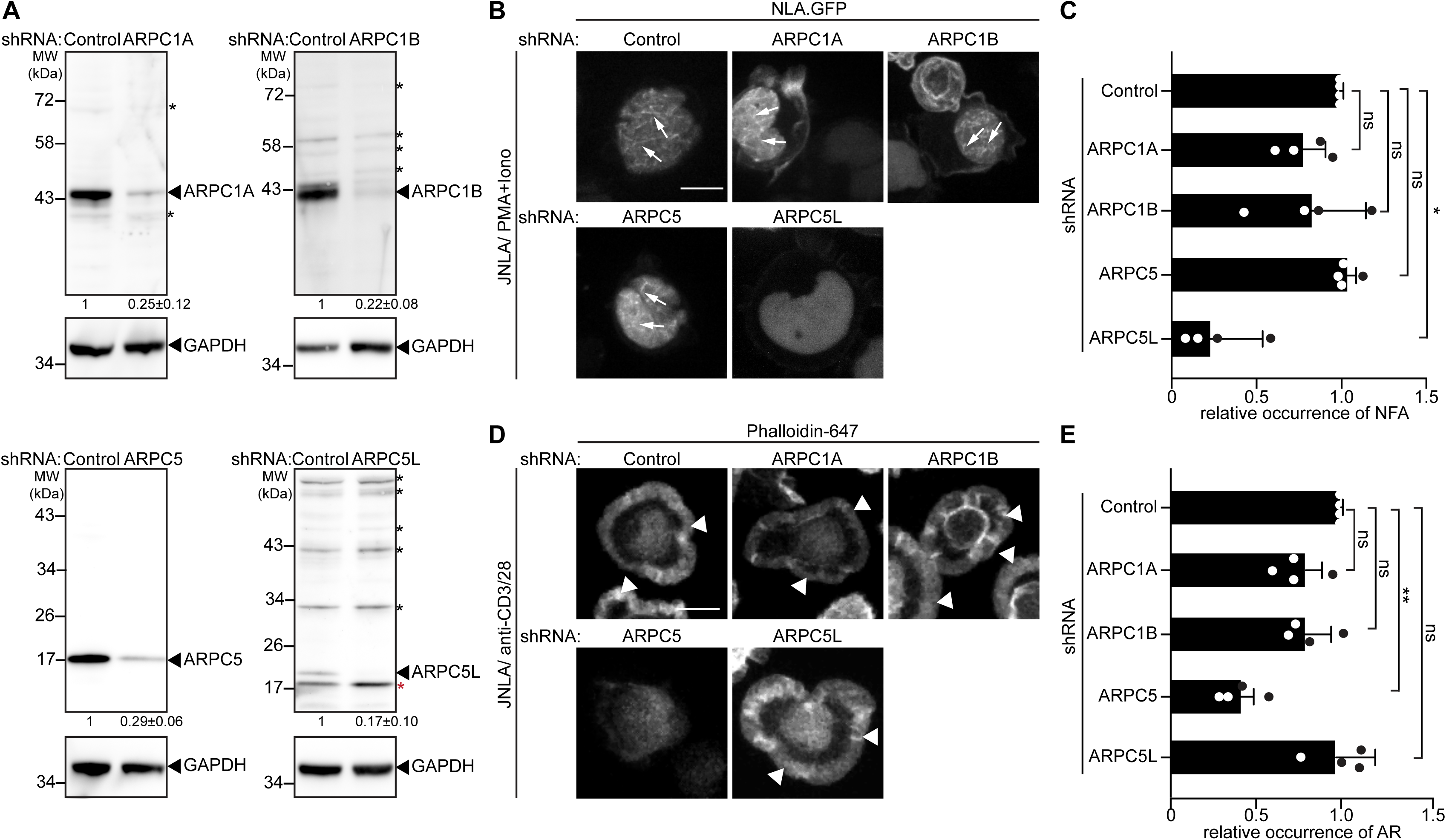
ARPC5 isoforms differentially regulate nuclear and plasma membrane actin polymerization. (**A**) Representative immunoblots show knockdown of ARPC1 and ARPC5 isoforms in bulk JNLA cells treated with indicated shRNA. Black arrowheads indicate the specific bands, black asterisks mark unspecific bands. Note that the ARPC5L antibody also detects ARPC5 (marked by red asterisk). The numbers indicated below the respective blots represent mean±S.D values from four independent experiments, based on the densitometric quantification of the bands, normalized to GAPDH & compared to the NTC protein levels (which is set to 1). (**B**) Representative spinning disk confocal still images of JNLA cells treated with indicated shRNA, show stills post activation with P/I. Arrows point to the nuclear F-actin (NFA). (**C**) Quantification of NFA formation in shRNA treated cells relative to the scrambled control treated cells. Error bars were calculated from mean±SD of 4 independent experiments where 30 cells were analyzed per condition per experiment. Each dot represents mean of each independent experiment. (**D**) Representative immunofluorescence images indicate averaged intensity projections of Phalloidin-647 stained F-actin ring (AR) formation in JNLA cells treated with indicated shRNA upon activation on coverslips coated with antiCD3+CD28 antibodies. Arrowheads point to the F-actin ring at the PM. (**E**) Quantification of Phalloidin stained F-actin ring (AR) formation in shRNA treated cells relative to the control treated cells. Error bars were calculated from mean±SD of four independent experiments where at least 100 cells were analyzed per condition per experiment. Each dot represents mean of each independent experiment. One-way ANOVA with Kruskal-Wallis test was used to determine statistical significances, where *\*P ≤ 0.0332, **P≤ 0.0021* and *ns*: not significant. Scale bar, 7µm.

### Knock-out and reconstitution of ARPC5 isoforms

Our observations pointed to a central role for ARPC5 and ARPC5L in the specificity of Arp2/3 driven cytoplasmic and nuclear actin polymerization in response to CD4 T cell activation. To validate our observations with transient silencing, we generated ARPC5 or ARPC5L knock out (KO) cell lines using CRISPR-Cas9 ribonucleoprotein transfection (Fig. 4). ARPC5 or ARPC5L levels were strongly reduced (by at least 80%) in the resulting bulk KO cultures as well as in clones expanded from single KO cells (Figs. S2A, S3A). The levels of other Arp2/3 subunits remained largely unaffected but ARPC1B expression was reduced by almost 2-fold compared to the non-targeting control (NTC) in both KO cultures and ARPC5 KO was associated with a reduction or increase of ARPC1A (to 76%) and ARPC5L (to 240%), respectively (Fig. S2A). The nucleofection and transduction procedures used to generate and study these KO cells slightly reduced the overall efficiency of NFA formation in response to T cell activation. Nevertheless, T cell stimulation confirmed that ARPC5 is selectively required for cytoplasmic actin polymerization and is not substituted by the elevated levels of ARPC5L. In turn, ARPC5L is essential for NFA formation and dispensable for cytoplasmic actin polymerization (Figs. 4A-B, see mCherry controls and Figs. S3B-E). Reintroduction of the respective mCherry tagged ARPC5 isoform in these bulk (Figs. 4A-B; S2B-C) and clonal (Figs. S3A-E) KO cells reconstituted their ability to form NFA or ARs, respectively, at expression levels that ranged from significantly lower levels than the endogenous protein to significant overexpression (Fig. S2B-C, S3A). ARPC5 and ARPC5L are thus involved in distinct Arp2/3-dependent actin polymerization events during CD4 T cell activation.

**Fig. 4.**
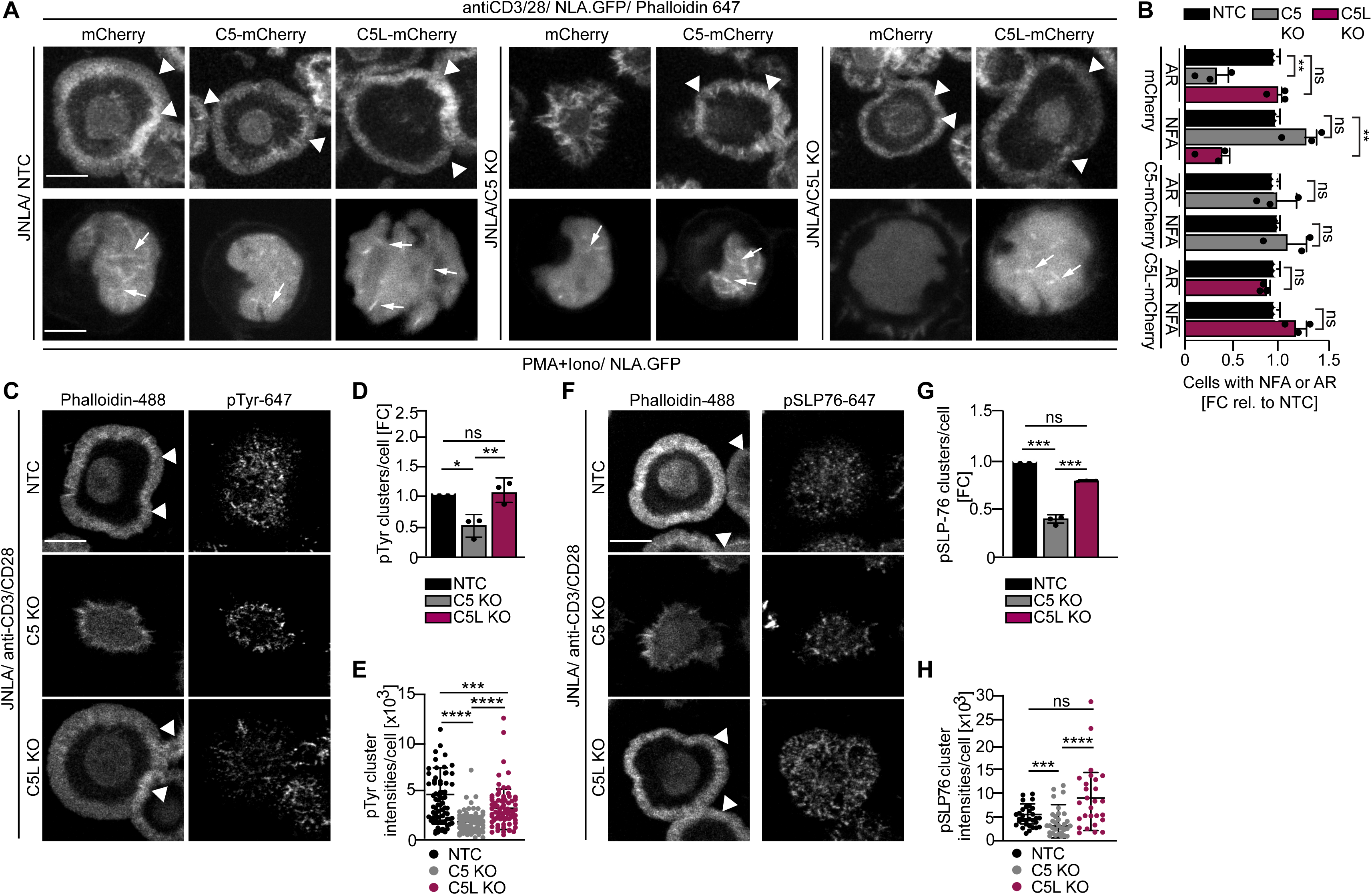
Effects observed on nuclear and plasma membrane F-actin dynamics upon ARPC5/C5L knockout and its impact on proximal TCR signaling. (**A**) Shown are representative maximum intensity projections of confocal still images of the indicated KO JNLA cells overexpressing mCherry (control) or mCherry fusion proteins of the respective ARPC5 isoforms, post activation with either anti CD3+CD28 antibodies (top panel) or with P/I (bottom panel). Arrows point to the nuclear F-actin (NFA, bottom). Arrowheads point to the F-actin ring (top). Scale bar, 7µm. **(B)** Quantification of AR formation in the PM, stained with Phalloidin647 is compared to the NFA formation visualized with NLA-GFP in the respective KO or KO+ARPC5 isoform expressing cells was performed relative to the non-targeting control (NTC) treated cells. ‘mCherry’ alone was used as vector backbone control for the overexpression study. Bars indicate mean from three independent experiment where 30-40 cells were analysed per condition. One-way ANOVA with Kruskal-Wallis test was used to determine statistical significances, where *\*\*P≤ 0.0021* and *ns*: not significant. **(C,F)** Representative confocal images of JNLA.GFP cells with indicated knockout or control (NTC), upon 5min of activation on coverslips coated with anti-CD3+CD28 antibodies. Cells were fixed and stained for F-actin (with Phalloidin 488) and pTyr or pSLP-76 (Alexa Fluor 647), respectively. **(D,G)** shows quantification of the total number of pTyr or pSLP-76 clusters/cell in KO relative to control cells analysed using the ‘Spot Detector’ Fiji plugin. **(E,H)** Dot plots represent the changes in overall intensity of pTyr or pSLP-76 clusters per cell where each dot represents intensity of clusters/cell analysed. Error bars were calculated from mean±SD of 3 independent experiments where ∼80-100 cells were analyzed per condition. One sample t-test was used to determine statistical significances, where *\*P* ≤ 0.033, *\*\*P* ≤ 0.0021, *\*\*\*P* ≤ 0.0002 and ns: not significant. Scale bar, 7µm.

### Nuclear actin dynamics is dispensable for TCR proximal signaling

Actin polymerization at sites of TCR engagement is directly coupled to downstream signalling constituted by dynamic phosphorylation cascades occurring in microclusters (Monks, Freiberg et al. 1998, Grakoui, Bromley et al. 1999, Dustin, Tseng et al. 2006). Nuclear actin dynamics precedes cytoplasmic actin polymerization upon TCR stimulation but whether NFA formation affects microcluster formation or function is unclear. We therefore tested whether ARPC5 isoforms differently impact generation and composition of these microclusters formed in response to surface-mediated TCR engagement. Disruption of cytoplasmic actin polymerization upon KO of ARPC5 significantly reduced the number of signalling microclusters as well as the amount of tyrosine phosphorylation (pTyr) or phospho-SLP-76 (pSLP-76) within the microclusters (Fig. 4C-E and 4F-H). In contrast, loss of ARPC5L had no effect on the amount of TCR signaling-induced microclusters formed or their pSLP-76 content. In turn, the pTyr intensity in these microclusters was reduced relative to that in control cells, albeit to significantly lower extend than in C5 KO cells. Together, these results revealed that TCR proximal signaling in response to CD4 T cell activation is governed by ARPC5-mediated actin polymerization at the plasma membrane and occurs independently from the formation of nuclear actin filaments mediated by ARPC5L containing Arp2/3 complexes.

### Subcellular localization and association with NFA does not determine the functional specificity of ARPC5 isoforms

We next assessed whether the differential role of ARPC5 and ARPC5L in TCR-induced actin remodeling reflects their distinct cellular distribution. Since antibody staining did not allow to distinguish between the distribution of endogenous ARPC5 and ARPC5L, we examined the localization of transiently expressed, mCherry-tagged isoforms that functionally rescued our KO cell lines. Both mCherry-tagged ARPC5 and ARPC5L had a diffuse cytoplasmic distribution but were also detected as punctae in the cytoplasm and the nucleus (Fig. 5A). Staining of endogenous ARPC5 and ARPC5L by the non-discriminating anti-ARPC5 antibody revealed similar punctae in the cytoplasm and nucleus with an additional localization at the plasma membrane that was particularly pronounced following surface-mediated TCR engagement (Figs. S4A-B). Consistently, immunoblot analysis of nucleo-cytoplasmic fractionations revealed that endogenous ARPC5 and ARPC5L are both present in the nucleus, albeit at lower levels than in the cytoplasm (Fig. 5B). This distribution of ARPC5 and ARPC5L was unaffected by the loss of expression of the other ARPC5 isoform. To assess the localization of ARPC5.mCherry and ARPC5L.mCherry relative to the NFA network, we applied two-color super resolution STED microscopy on P/I-stimulated A301 CD4 T cells, which are best suited to visualize endogenous NFA meshwork in T cells (Tsopoulidis, Kaw et al. 2019). Deconvolved and segmented STED images revealed a complex NFA meshwork (Fig. 5C). ARPC5.mCherry and ARPC5L.mCherry were both detected in discrete spots within the nucleus and ∼ 10% of these spots co-localized with nuclear actin filaments, however, no significant difference was observed between both isoforms (9.9% for ARPC5, 13.2% for ARPC5L, Fig. 5D). The identity of the ARPC5 isoform involved therefore does not determine the ability of Arp2/3 complexes to associate with actin filaments in the nucleus.

**Fig. 5.**
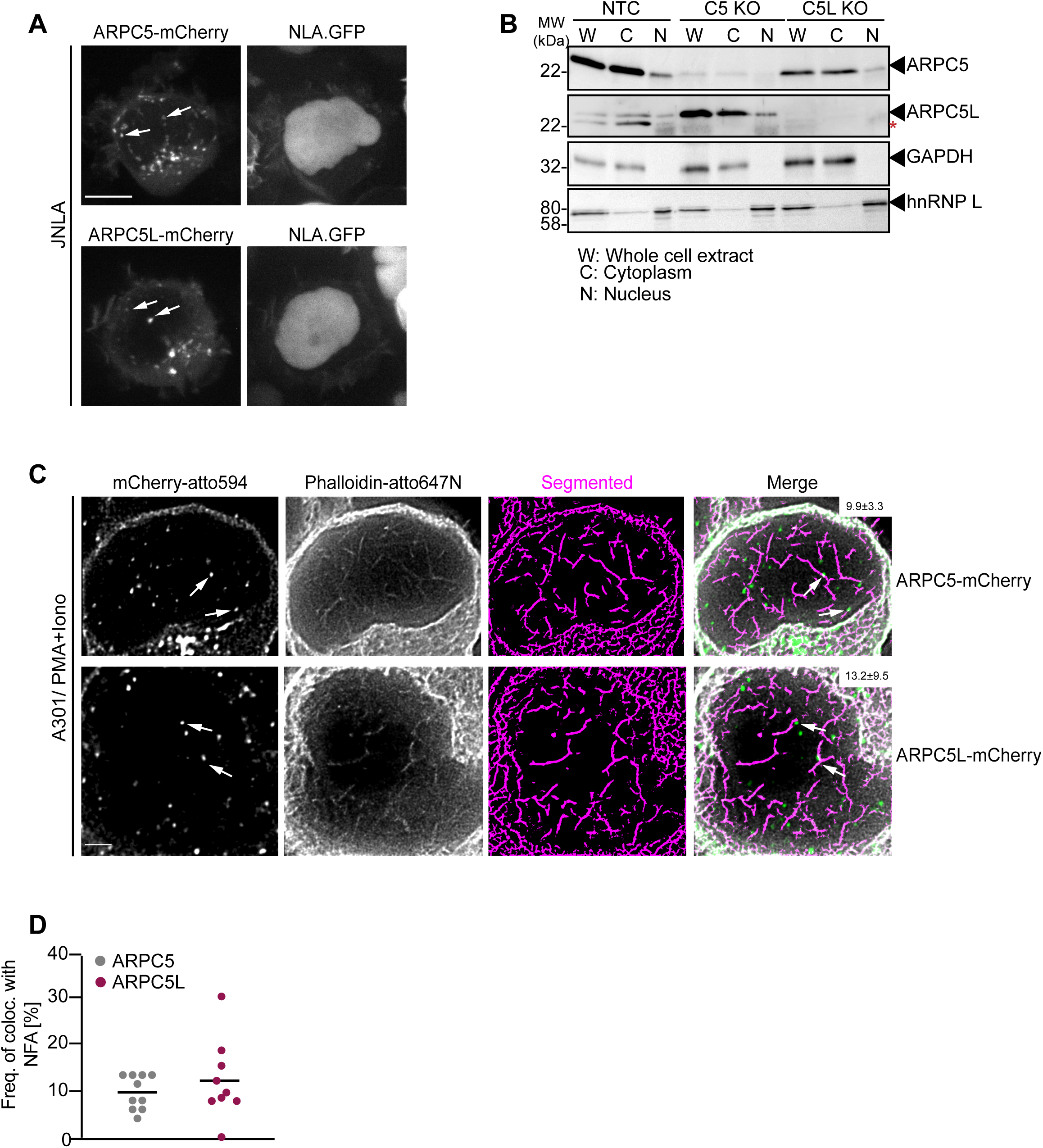
Cellular distribution of ARPC5 isoforms. **(A)** Shown are representative spinning disk confocal images (maximum projection) of ARPC5.mCherry and ARPC5L.mCherry distribution in unstimulated JNLA cells. White arrows point to the respective C5 or C5L punctae seen in the nucleus. **(B)** Subcellular distribution of the ARPC5 subunit and its isoform ARPC5L was determined by biochemical fractionation of JNLA cells with respective knockout of either C5 or C5L in the bulk culture. Representative immunoblots reveal levels of ARPC5 isoforms in the whole cell extract (WCE), cytoplasmic (C) and nuclear (N) fractions in the indicated JNLA knockout cells post fractionation. GAPDH and hnRNPL were used as markers for cytoplasmic and nuclear compartments, respectively. (**C**) Representative, deconvoluted and segmented Stimulated emission depletion (STED) single plane images show endogenous nuclear actin filaments (stained with Phalloidin-647N) and ARPC5.mCherry/ARPC5L.mCherry (signal enhanced with anti-mCherry with secondary antibody in atto-594 channel) in A3.01 T cells, stimulated with P/I for 30s. Arrows (in white) point to the colocalization events. Percent colocalization is mentioned as mean± SD (in white bar, top right) for each of the isoforms from three independent experiments. Scale bar, 500nm. **(D)** The dot plot shows frequency of colocalization of ARPC5 and ARPC5L with nuclear F-actin (from representative STED-deconvolved & segmented super-resolved images shown in Fig3D) in A3.01 cells post 30s of stimulation with PMA+Ionomycin. Each dot represents colocalization events per cell that was analysed.

### NFA formation triggered by DNA replication stress involves ARPC5 but not ARPC5L and is not regulated by nuclear calcium transients

We next tested whether the selective involvement of ARPC5L is a common principle for Arp2/3-dependent nuclear actin polymerization events. In fibroblasts and epithelial cells, Arp2/3 also mediates nuclear actin polymerization in response to DNA replication stress induced by the DNA polymerase inhibitor Aphidicolin (APH) (Lamm, Read et al. 2020). Consistent with that fact that effective drug concentrations are often elevated in CD4 T cell lines (Martel, Payet et al. 1997, Vesela, Chroma et al. 2017), we observed efficient formation of NFA in JNLA cells in response to APH starting at 15µM concentration (Fig. S5A). This response was paralleled by the induction of DNA replication stress as indicated by phosphorylation of the checkpoint kinase CHK-1 (Fig. S5B). NFA formation in response to APH was transient with a maximum of cells displaying NFA approx. 90 min post treatment (Figs. 6A-C, see Fig. S5C for tracks of individual cells). This APH-induced NFA meshwork appeared to consist of fewer but thicker F-actin bundles and to disassemble more slowly than NFA induced by T cell activation (Fig. 6B). Next, ARPC5L and ARPC5 KO cells were stimulated in parallel by P/I or APH. For T cell activation, this confirmed the requirement of ARPC5 for formation of actin rings (AR) and ARPC5L for the NFA network (Fig. S5D). In contrast, NFA formation in response to APH was indistinguishable to control cells in ARPC5L KO cells but significantly impaired in ARPC5 KO cells (Figs 6B, D). NFA induction by T cell activation or DNA replication stress is thus mediated by specific Arp2/3 complexes containing distinct ARPC5 subunit isoforms and nuclear localization of the actin polymerization event alone does not govern the involvement of the ARPC5 or ARPC5L isoforms. We therefore tested if this specificity for ARPC5 isoforms is provided by upstream signaling. NFA formation induced by T cell activation is mediated by nuclear calcium transients and can be inhibited by interfering with nuclear calmodulin by expressing a dominant negative version of calmodulin binding protein 4 (CAMBP4) (Monaco, Jahraus et al. 2016, Tsopoulidis, Kaw et al. 2019) (Fig. 6E, F and Fig. S5E, F). In contrast, dominant negative CAMBP4 did not affect NFA formation or CHK-1 phosphorylation upon APH treatment of JNLA cells (Fig. 6E-F and S5B). Similarly, pharmacological inhibitors of downstream effectors of calmodulin including Calcineurin inhibitor CyclosporinA (CsA), Calmodulin-kinase kinase inhibitor STO609 as well as the Calmodulin-kinase II inhibitors KN93 and KN62 did not prevent NFA induction by APH (Fig. S5G). These results suggest nuclear Ca^2+^-calmodulin acts as a selective trigger for ARPC5L-dependent nuclear actin polymerization.

**Fig. 6.**
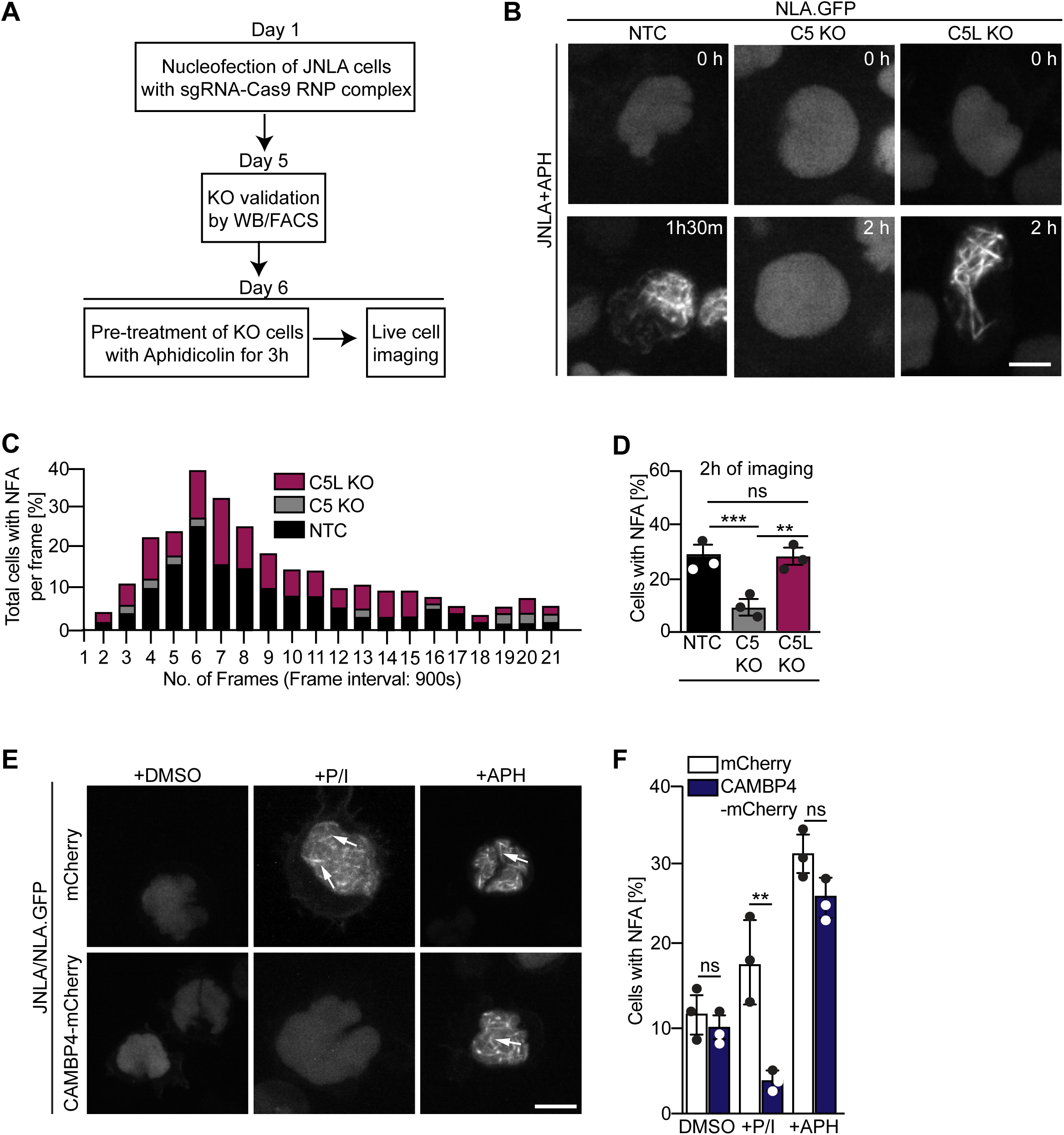
Differential role of ARPC5 isoforms in replication stress mediated NFA formation. **(A)** Schematic of experimental setup showing timeline of knockout (KO) generation and induction of replication stress in the JNLA KO cells using Aphidicolin (APH). **(B)** Shown are representative spinning disk confocal still images (maximum projection) of the APH pre-treated KO or control cells. The movies were acquired for 5h with acquisition every 15min post pre-treatment of cells with APH. The stills at the indicated time points are representative of the timepoint where the maximal NFA burst has been observed in each condition. **(C)** Stacked bar graph (denoted by three different colours for each condition) shows the kinetics of NFA burst throughout 5h of live cell imaging duration, with maximum NFA burst observed within the first 2h. **(D)** Quantification of the % of cells with NFA bursts (within first 2h of imaging) post replication stress induction in control and KO cells are shown where the bars represent mean values from three independent experiments. Around 40-60 cells/condition/experiment where analyzed. Statistical significance was calculated using One-way ANOVA (Kruskal-Wallis test). **(E)** JNLA cells transduced with either mCherry (control) or CAMBP4.NLS-mCherry were pre-treated with solvent control (DMSO), were either activated by PMA+Ionomycin (P/I) or treated with APH for induction of replication stress for 3h prior to live cell imaging. Shown are maximum projection of representative spinning disk confocal still images (showing the timeframe where maximum NFA burst was observed) in the DMSO control compared with either P/I or APH mediated NFA bursts (white arrows) in the presence and absence of nuclear Calmodulin. Movies for visualizing replication stress were acquired for 5h with acquisition every 15min post pre-treatment of cells. Whereas movies for visualizing P/I activation induced NFA were acquired for 5mins with acquisition every 15-30s. **(F)** Quantification of NFA in the above-mentioned conditions were performed as shown here, are the bars representing mean values from three independent experiments, where at least 30 cells were analyzed per condition per experiment for NFA quantification. Statistical significance was calculated using Welch’s t-test. \**P* ≤ 0.0332, \*\**P* ≤ 0.0021, *\*\*\*P ≤ 0.0002* and ns: not significant. Scale bar, 5µm.

### The nucleation promoting factor N-WASP selectively drives TCR-induced NFA formation

Since the specificity of distinct nuclear actin polymerization events for distinct ARPC5 isoforms is determined upstream by nuclear Ca^2+^ transients, we assessed if specific NPFs are involved in this process and focused on class I NPFs with reported roles in the nucleus (Teitell Michael 2010, Weston, Coutts et al. 2012, Wang, Du et al. 2022). While we were unable to generate WASp KO JNLA cells, KO of N-WASP, WASHC5, an essential subunit of the WASH regulatory complex that is required for the NPF function of WASH (Jia, Gomez et al. 2010), or WAVE2 resulted in bulk cultures with significantly reduced expression levels (Fig. 7A). Functional characterization revealed the specific involvement of these NPFs in distinct actin polymerization events: while N-WASP was essential for efficient NFA formation in response to P/I, WASHC5 was dispensable for NFA formation and loss of WAVE2 did not affect the frequency of NFA formation but was associated with the formation of shorter nuclear actin filaments (Figs. 7B, C). In contrast, N-WASP was dispensable for cell spreading and AR formation induced by TCR signaling while loss of WASHC5 or WAVE2 significantly impaired these processes (Fig. 7D, E, S6D). Interestingly, NFA formation in response to APH was unaffected in N-WASP, WASHC5 or WAVE2 KO cells (Fig. 7F, G) and is hence induced by other upstream regulators. NFA formation by TCR signaling is thus governed by a selective pathway that depends on ARPC5L containing Arp2/3 complexes that are regulated by N-WASP and nuclear Ca^2+^ transients. Interestingly, while expression levels of Arp2/3 subunits and isoforms were overall unaltered in NPF KO cells, ARPC5L levels were significantly reduced in N-WASP KO cells (Fig. S6A-C). Functional coupling of ARPC5L and N-WASP may thus involve a regulatory mechanism at the level of protein expression/stability.

**Fig. 7.**
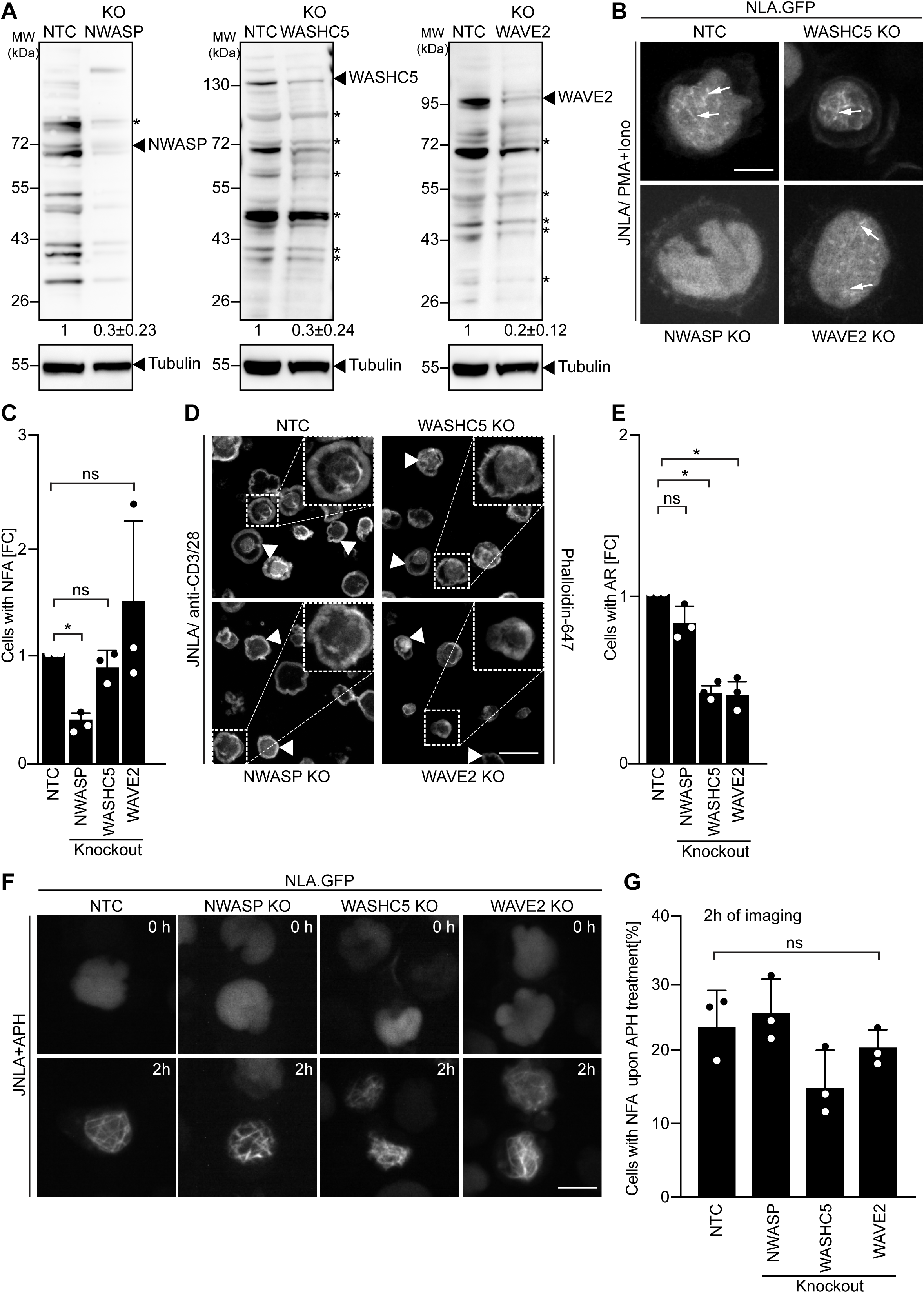
Differential involvement of class I NPFs in T cell activation and replication stress mediated NFA formation. (**A**) Representative immunoblots show knockout (KO) of NWASP, WASHC5 and WAVE2 class I NPFs, respectively, in JNLA cells. Black arrowheads indicate the specific bands, black asterisks mark unspecific bands. The numbers indicated below the respective blots represent mean±S.D values from three independent experiments, based on the densitometric quantification of the bands, normalized to Tubulin & compared to the NTC protein levels (which is set to 1). (**B**) Representative spinning disk confocal still images (maximum projection) of JNLA cells with indicated NPF KO, showing NFA formation post activation with P/I. Movies were acquired for 5min post PI addition with acquisition every 30s. Arrows point to the nuclear F-actin (NFA). (**C**) Quantification of NFA formation in respective NPF KO cells relative to the non-targeting control (NTC) treated cells. Error bars were calculated from mean±s.d of three independent experiments where 30 cells were analysed per condition per experiment. Each dot represents mean of one independent experiment. (**D**) Representative confocal images indicate averaged intensity projections of Phalloidin-647 stained F-actin ring (AR) formation in fixed/permeabilized JNLA cells with respective NPF KO upon activation on coverslips coated with antiCD3+CD28 antibodies. Arrowheads point to the f-actin ring at the PM upon TCR activation and the dotted box in white shows the cell in zoomed view in the inset (top right). (**E**) Quantification of Phalloidin stained F-actin ring (AR) formation in KO cells relative to the NTC treated cells. Error bars were calculated from mean±s.d of three independent experiments where at least 100 cells were analysed per condition per experiment. Each dot represents mean of one independent experiment. **(F)** Shown are representative spinning disk confocal still images (maximum projection) of the APH treated NPF KO or control cells, respectively. The movies were acquired for 5h with acquisition every 15min post pre-treatment of cells with APH. The stills at the indicated time points are representative of the timepoint where the maximal NFA burst has been observed in each condition. **(G)** Quantification of the % of cells with NFA bursts (within first 2h of imaging) post replication stress induction in control and KO cells are shown where the bars represent mean values from three independent experiments. Around 40-60 cells/condition/experiment where analysed. One-way ANOVA with Kruskal-Wallis test was used to determine statistical significances, where *\*P ≤ 0.0332* and *ns*: not significant. Scale bar, 5µm

## Discussion

A main finding of our study is that nuclear and cytoplasmic actin polymerization triggered by a common stimulus (T cell activation) or nuclear actin polymerization by different stimuli (T cell activation, DNA replication stress) are mediated by distinct Arp2/3 complexes containing ARPC5L or ARPC5, respectively. Since (i) nuclear actin polymerization events induced by TCR signaling or APH-mediated induction of DNA replication stress were also mediated by Arp2/3 complexes with distinct ARPC5 subunit preferences and (ii) Arp2/3 complexes containing both isoforms are present and operational in the nucleus, the selectivity for an ARPC5 isoform is not determined at the level of subcellular distribution. Rather, the nature of the stimulus is critical for the selective induction of actin polymerization by ARPC5 or ARPC5L containing Arp2/3 complexes (see schematic model in Fig. 8). Our results define responsiveness to nuclear calcium-calmodulin signaling and regulation by the NPF N-WASP as specific triggers of ARPC5L containing complexes. The selective involvement of N-WASP likely reflects that its ability to activate Arp2/3 can be achieved by calcium-calmodulin, directly or via activation of its regulator IQGAP (Miki, Miura et al. 1996, Le Clainche, Schlaepfer et al. 2007, Pelikan-Conchaudron, Le Clainche et al. 2011). How ARPC5L determines the sensitivity of Arp2/3 to regulation by N-WASP-calcium-calmodulin at the molecular level remains to be determined. In principle, the identity of the ARPC5 isoform may affect the affinity of Arp2/3 to activation by N-WASP. However, the available structural and biochemical insight does not support a direct role of ARPC5/C5L in interactions with NPFs (von Loeffelholz, Purkiss et al. 2020). ARPC5 isoforms thus likely undergo specific interactions with additional interaction partners that govern the susceptibility of Arp2/3 complex to activation via N-Wasp-calcium-calmodulin. Indeed, a recent preprint by Fäßler and colleagues reports that in migrating fibroblasts, the identity of the involved ARPC5 isoform affects the positioning of the Arp2/3 effectors Mena/VASP and thereby filament polymerization velocity (Faessler, Javoor et al. 2022). In addition to defining this mechanism it will be interesting to determine whether induction of ARPC5L-containing complexes by calcium-calmodulin can also occur in the cytoplasm and how the observed stabilization of ARPC5L expression by N-WASP contributes to the regulation of this pathway. In contrast, Arp2/3 complexes containing ARPC5 such as those involved in DNA replication stress trigger nuclear actin polymerization independently of calcium-calmodulin-N-WASP and are likely regulated by another NPF. It is tempting to speculate that the regulation of ARPC5L containing Arp2/3 complexes by nuclear calcium-calmodulin reflects the requirement for rapid conversion of an extracellular signal, e.g., to elicit a transcriptional response. In line with this scenario, calcium-mediated induction of nuclear actin assembly by the formin INF2 in mouse fibroblasts also represents a rapid response to conversion of an extracellular signal (Wang, Sherrard et al. 2019). In contrast, DNA replication stress provides a signal from within the nucleus without the need for a fast second messenger. Notably, T cell activation or DNA replication stress induce NFA networks of different filament morphology and dynamics. These architectural differences may reflect physiological properties of Arp2/3 complexes with different ARPC5 isoforms and translate into distinct functional roles. In this scenario, thin/dynamic filaments may mediate transcriptional regulation following TCR engagement while thicker and more stable filaments could exert mechanical functions during DNA repair. The preference for distinct ARPC5 isoforms thus likely adjusts the activity of Arp2/3 complex to such divergent actin polymerization events that are triggered by specific upstream signals. Similarly, Arp2/3 complexes containing different ARP3 isoforms were recently shown to be differentially regulated (Abella, Galloni et al. 2016, Galloni, Carra et al. 2021). The subunit isoform composition of Arp2/3 complexes thus emerges as important parameter that allows Arp2/3 to mediate distinct actin polymerization events tailored to specific activation signals at selected subcellular sites.

**Fig 8.**
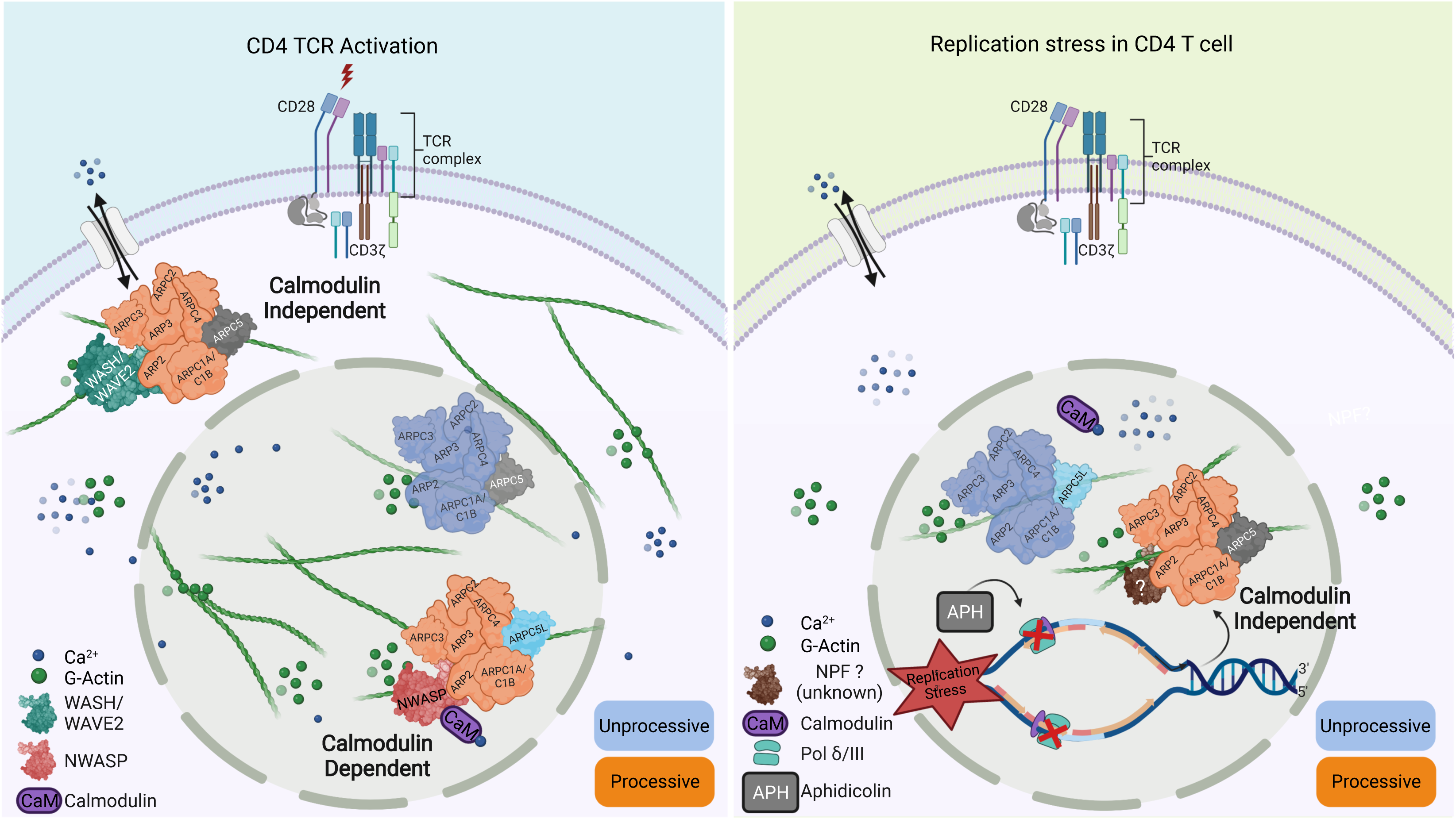
Graphical summary of our findings. Schematic model for Arp2/3 dependent differential regulation of actin dynamics induced upon TCR engagement (left) when compared to induction of replication stress by aphidicolin (APH) (right).

## Acknowledgments

We are grateful to Nadine Tibroni and Ina Ambiel for technical assistance and Kathrin Bajak for help with manuscript preparation and submission. This project is supported by the Deutsche Forschungsgemeinschaft (DFG, German Research Foundation) by project FA 378/20-1 to OTF. MW was supported by Cancer Research UK (FC001209), the UK Medical Research Council (FC001209), and the Wellcome Trust (FC001209) funding at the Francis Crick Institute as well as by the European Research Council (ERC) under the European Union’s Horizon 2020 research and innovation programme (grant agreement No 810207 to MW). For the purpose of Open Access, the author has applied a CC BY public copyright licence to any Author Accepted Manuscript version arising from this submission.

## Author contributions

Conceptualization, O.T.F., N.T., Methodology, L.S., N.T., V.L.; Investigation, L.S., N.T., V.L.; Data analysis, L.S., N.T., V.L.; Writing – Original Draft, O.T.F., L.S.; Writing – Review & Editing, O.T.F, M.W., L.S., N.T., V.L.; Funding Acquisition, O.T.F.; Resources, M.W; Supervision, O.T.F

## Materials and Methods

### Cells and reagents

HEK 293T cells were cultured in DMEM high glucose plus 10% feline bovine serum (FBS, Milipore), 100U/mL penicillin and 100μg/mL streptomycin. Primary T cells, Jurkat Tag cells (JTAgs) and CLEM derived A3.01 cells were cultured in RPMI containing 10% FBS and 1% Penicillin-streptomycin and GlutaMAX-I (Gibco). All experiments performed in JTag cells stably expressing nuclear lifeact-GFP (JNLA) were obtained as described previously in (Tsopoulidis, Kaw et al. 2019). All Cell lines were cultivated according to their ATCC (https://www.atcc.org) guidelines. For visualization of nuclear F-actin, A3.01 or JNLA were washed thoroughly with PBS, adjusted to a cell density of 3E5/ml, and incubated overnight in RPMI (phenol-Red free medium, GIBCO) containing 0.5% (A3.01) or 10% (JNLA) FBS. For Immunofluorescence (IF) microscopy: F-actin was stained with Phalloidin Alexa Fluor 488 or atto-AF488 (Thermo Fischer). Alexa antibodies for IF such as: goat anti-mouse Alexa Fluor 568, goat anti-rabbit Alexa Fluor 647 and goat anti-rabbit Alexa Fluor 568 were obtained from Thermo Fischer Scientific. The following anti-CD3 (clone HIT3a against CD3ε; BD Pharmingen) and mouse anti-CD28 (CD28.2, BD Pharmingen) were used at 1:100 dilution for coating coverslips/GBDs to make stimulatory surface for T cell activation. Antibodies used for immunoblotting were mouse-anti-ARP3, 1:10,000 (cloneFMS338, SIGMA), mouse anti-p16-ARC, 1:500 (#305011, Synaptic systems), rabbit anti-ARPC5L, 1:1000 (GTX120725 GeneTex), rabbit anti-ARPC1A, 1:500 (#HPA004334, Sigma), mouse anti-ARPC1B, 1:500 (SCBT), rabbit anti-ARPC2, 1:1000 (EPR8533 Abcam), mouse anti-ARPC3, 1:500 (#HPA006550, Sigma Aldrich), mouse anti-ARPC4, 1:500 (#NBP1-69003, Novus Biologicals), mouse anti-WAVE2, 1:500 (#sc-373889, SCBT), rabbit anti-NWASP/WASL, 1:1000 (#HPA005750, Sigma Aldrich), rabbit anti-WASHC5, 1:250 (#HPA070916, Sigma Aldrich), mouse anti-Tubulin, 1:1000 (#373, DM1A, CST), rabbit anti-GAPDH, 1:2500 (#2118, 14C10, CST), mouse anti-GAPDH, 1:2000 (#MCA4740, BioRad), mouse anti-mCherry, 1:1000 for WB and 1:500 for IF (NBP1-96752), rabbit anti-mCherry, 1:1000 for WB and 1:500 for IF (ab167453), rabbit anti-pTyr, 1:100 (#sc18182, SCBT), rabbit anti-pSLP76, 1:1000 (#ab75829, Abcam), HRP-coupled secondary rabbit or mouse antibodies for immunoblotting were obtained from Jackson Immuno Research was used at a dilution of 1:5000 for all samples. The secondary Alexa fluorescent coupled antibodies (either mouse or rabbit) used for IF staining were obtained from Invitrogen and were used at a dilution of 1:1000. For live cell imaging and STED microscopy: glass-bottom-dishes (GBD) with 35 mm plate diameter, 14 mm glass diameter, thickness 1.5 (Mattek corporation) and µ-slide 8-well glass bottom chambers (Ibidi) were used along with poly-lysine (Sigma), coated at a concentration of 0.01% in sterile filtered water. Phalloidin atto-647N used for STED imaging was bought from ATTO-TEC GmbH (AD 647N-81) and used at a dilution of 1:500 in blocking buffer containing 3% FCS/PBS for immunofluorescence staining whereas for super resolved STED imaging Phalloidin was dissolved in 5% FCS/cytoskeleton buffer (1X).

### Preparation of primary CD4 T cells

For the isolation of primary human CD4 T cells, human Buffy Coats from anonymous healthy donors were obtained from the Heidelberg University Hospital Blood Bank. CD4+ T cells were isolated by negative selection with the RosetteSepTM Human CD4+ T Cell Enrichment Cocktail and separated by Ficoll gradient centrifugation, resulting in homogenous populations of CD4+ T cells with a purity of 90-95% as assured by flow cytometry. Cells labelled as ‘Resting’ were cultured for 72h in complete RPMI media containing recombinant human IL2 (Biomol #155400.10) at 10ng/ml final concentration. Whereas the cells labelled as ‘Activated’ were cultured for 72h in complete RPMI media containing recombinant human IL2 (Biomol #155400.10) at 10ng/ml final concentration along with dynabeads at a ratio of 25µl Human anti CD3/28 labelled Dynabeads/10 million cells (#11132D, Gibco).

### Agonists and inhibitors

The following chemicals were used at the indicated concentrations: Ionomycin (Iono, 2μM), Phorbol 12-myristate 13-acetate (PMA, 162 nM), CK-869 (100 μM), KN-93 (0.25 μM), KN-62 (2.5 μM), STO-609 (5μM), and Aphidicolin (15µM) were all obtained from Sigma Aldrich whereas Cyclosporine A (1µM) was obtained from TOCRIS bioscience.

### Expression plasmids

pLVX vector expressing either human ARPC5/C5L cDNA fused to a mCherry fluorescent reporter or just the mCherry alone, were a kind gift from the lab of M.Way, generated as described in (Abella, Galloni et al. 2016). Plasmids expressing mCherry alone or mCherry conjugated to CAMBP4 in the pWPI backbone were used as described (Tsopoulidis, Kaw et al. 2019) and were selected using blasticidin (5µg/ml) for 48-72h. For the stable expression of shRNAs, gene specific target sequences (available upon request) were cloned into the lentiviral vector pLKO.1-puro (Addgene) as described in (Tsopoulidis, Kaw et al. 2019) and were selected with Puromycin (1.5 µg/ml) for 48h post transduction.

### Live-cell imaging of actin dynamics

Live imaging of actin dynamics was performed with a Nikon Ti PerkinElmer UltraVIEW VoX spinning disc confocal microscope equipped with a perfect focus system (PFS), a 60X oil objective (numerical aperture, 1.49), Hamamatsu ORCA-flash 4.0 scientific complementary metal-oxide semiconductor camera, and an environmental control chamber (37°C, 5% CO2), as described earlier in (Tsopoulidis, Kaw et al. 2019). Acquisition settings varied depending on the total acquisition time required for each experimental question/setup. The following are the acquisition settings for short term imaging: exposure time, 300 ms; frame rate, 6 to 10 frames/s, number of Z planes, 10; Z-stack spacing, 0.5µm; 488 nm, laser power between 4.5-5.5%; and total acquisition time, 3 to 10 min. Jurkat cells stably expressing nuclear lifeact.GFP (JNLA) were always washed with PBS and split 24 hours before the experiment to a density of 3×10^5^/ml. The next day, 3×10^5^ cells were harvested, washed with PBS, and resuspended in 100µl reconstituted RPMI containing 10% FCS. For imaging the actin dynamics of JNLA cells falling on stimulatory surface (coated with anti-CD3+CD28 antibodies), the PFS system was adjusted first with a low amount of highly diluted cells placed on the coated glass bottom dish. A single cell was centered to the field of view, and the PFS was adjusted to automatically focus on the glass-cell contact site. Subsequently, the stage was moved to a cell-free area, and 100µl of the cell suspension (3 × 10^5^/100µl) was added to the glass bottom dish with simultaneously recording cells while making contact with the glass surface.

For imaging of actin dynamics in cells resting on the glass surface, 100µl of cells (3 × 10^5^/ml) was plated on polyK-coated glass-bottom dishes, allowed to adhere for 5 min, and then stimulated with PMA/Iono in RPMI media.

### Super resolution imaging of nuclear actin

A3.01 T lymphoblastoid cells were washed and adjusted to cell density of 0.35 million cells/ml a day prior to the experiment, as described above. The next day, 0.6 million cells were collected/well of a 8-well chambered dish, washed once with PBS and resuspended in of RPMI (phenol-free) containing 0.5% FBS. Cells were allowed to adhere for 5 min on polyK-coated 8-well chamber glass bottom dish. Stimulation was performed by adding 100 µl of PMA/Iono solution dropwise to the cell suspension. Cells were activated for 30s and then permeabilized and stained with 100µl Perm solution containing 0.3% Triton X-100 + Phalloidin -Alexa Fluor 488 (1:2000) in 1X cytoskeleton buffer [10 mM MES, 138 mM KCl, 3 mM MgCl, 2 mM EGTA, and 0.32 M sucrose (pH 6.1)] for 30s. Addition of low amounts of phalloidin at this step is required to stabilize nuclear actin filaments during permeabilization/fixation. Cells were fixed with 1 ml of 4% methanol-free formaldehyde (Pierce) in 1X cytoskeleton buffer and incubated for 25 min at RT in the dark. Subsequently, the fixed cells were washed twice with cytoskeleton buffer, blocked with 5% bovine serum albumin (BSA) prepared in 1X cytoskeleton buffer, and stained with 1:500 Phalloidin atto-647N in 1X cytoskeleton buffer for 1 hour at room temperature (RT) or overnight (ON) at 4°C. Additionally, to enhance the mCherry signal of the C5/C5L-mCherry expression constructs in the cells, primary antibody staining using anti-mCherry antibody (1:500) was performed ON in blocking buffer as mentioned above. This was followed by multiple washing steps in 1X cytoskeleton buffer and staining with secondary antibody conjugated with an Atto-568 dye, for 1h at RT. Phalloidin atto-647N was added at this step with the secondary antibody to stain the endogenous nuclear actin filaments, which were observed in ∼40-60% of the cells.

STED microscopy was performed on an Expert Line STED system (Abberior Instruments GmbH, Göttingen, Germany) equipped with an SLM-based easy3D module and an Olympus IX83 microscope body, using a 100x oil immersion objective (NA, 1.4; Olympus UPlanSApo). STED images were acquired using the 590 nm (ARPC5/ARPC5L signals) and 640 nm (actin filament signals) excitation laser lines in the line sequential mode with corresponding 615/20 and 685/70 emission filters placed in front of avalanche photodiodes for detection. 775 nm STED laser (15% of the maximal power of 3 mW) was used for depletion with pixel dwell time of 10 to 15 μs, 15 nm xy sampling and 9x accumulation. To increase the signal-to-noise and facilitate subsequent image segmentation and quantification, STED images were restored with Huygens Deconvolution (Scientific Volume Imaging) using Classic Maximum Likelihood Estimation (CMLE) algorithm and Deconvolution Express mode with “Conservative” settings. To segment actin filaments and ARPC5/ARPC5L signals in obtained STED images we trained a Random Forest classifier using ilastik (ref PMID: 31570887) autocontext workflow which predicts semantic class attribution (signal or background) for every pixel. The training set of data was arbitrary selected and very sparsely labelled (<0.1% of total pixels were manually categorized into “signal” and “background” categories). Obtained machine learning algorithm was used applied to all acquired images ensuring an unbiased signal segmentation across all experiments. This allowed the quantification of the number of ARPC5/ARPC5L signals colocalizing with nuclear actin filaments by visual inspection of binary (segmented) images.

### Imaging actin dynamics at the Immune synapse post CK-inhibitor treatment

To distinguish B cells from T cells before mixing them together for live imaging, Raji B cells were stained with Cell trace Deep Red (10μM, Thermo Fischer) at 1:1000 dilution for 1h and simultaneously loaded with Staphylococcal enterotoxin E (SEE, Toxin Technology) at a concentration of 5 μg/ml, in RPMI complete media for 30 min at 37°C and subsequently washed and resuspended in 10% FBS containing RPMI at a concentration of 5x10^4^ cells in 100μl. JNLA cells were washed and adjusted a day before as described above. 24h later 1x10^6^ were harvested, washed in PBS and resuspended in 100µl RPMI complete media containing either DMSO or the CK869 for 1h at 37°C. The media is replenished after 1h with fresh media containing either the solvent or the inhibitor such that the cells are at a final density of 5x10^4^ cells in 100μl. 100µl of the treated JNLA cells are plated on a poly-lysine coated GBDs. Approx. 5-10 regions on the GBDs were selected for live cell imaging using the spinning disk confocal microscopy as described above. Imaging was started and 100μl Raji B cells were added dropwise onto the T cell suspension while the image acquisition was ongoing. The following are the acquisition settings for the imaging: exposure time, 200-300 ms; frame rate, 6 to 10 frames/s, number of Z planes, 3; Z-stack spacing: 1-1.5µm; 488 nm, laser power 5.5%; and total acquisition time, 30min with acquisition every 30s/XY position.

### RNA extraction and Quantitative PCR (qPCR)

For RNA extraction NucleoSpin RNA II kit (Macherey-Nagel) was used. 10x10^6^ cells were collected per condition/per cell line, washed with cold PBS once and their pellets were stored at -80°C for maximum 2-3weeks. RNA extraction was done following manufacturer’s protocol. After RNA quantification by UV/VIS spectrometry (Nanodrop), between 500ng-1000ng of total RNA was reverse transcribed using the SuperScriptII (Invitrogen) according to the manufacturers’ instructions. 1:10 dilution of the cDNA in RNAse free water was used for qPCR reaction using the SYBR green PCR master mix (Life Technologies), and reactions were performed on a Quant Studio1 sequence detection system (Applied Biosystems) using the following program: 50°C for 2 min, 95°C for 10 min, and 40 cycles of 95°C for 15 s and 60°C for 1 min. GAPDH (glyceraldehyde-3-phosphate dehydrogenase) mRNA was used for normalization of input RNA wherever needed or mentioned. The primers used are available upon request.

### Immunoblot analysis

1xE6 cells/condition were collected and lysed in lysis buffer (50 mM Tris-HCl [pH 7.4], 75 mM NaCl, 1 mM EDTA, 1 mM NaF, and 0.5% NP-40) with a freshly added protease inhibitor cocktail and sodium vanadate and subjected to 9 cycles (30s ON-10s OFF) of ultrasonication (Bioruptor Plus; Diagenode). The sonicated samples are then spun down, and the supernatant is collected for protein estimation using the microBCA kit (Pierce). 10µg of protein is then mixed with 1X sample buffer (10% sucrose, 0.1% bromophenol blue, 5 mM EDTA [pH 8.0], 200 mM Tris [pH 8.8]), and boiled at 95°C for 10min. The lysates are then run on either self-made 10-15% SDS-PAGE gel or on pre-casted Invitrogen™ NuPAGE™ 4 bis 12 %, Bis-Tris, 1,0–1,5 mm, Mini-Protein-Gel, followed by blotting with Trans-blot PVDF membranes (BioRad) for 15min, blocked in 5% BSA in TBS-T for 1h before probing with the primary antibodies overnight at 4℃. Secondary antibodies conjugated to HRP were probed for 1-1.5h at room temperature the next day following 3x intensive TBST washing of the unbound primary antibodies. Enhanced chemiluminescence (ECL)-based detection using the WesternBright Sirius Chemiluminescent Detection Kit (Advansta) was performed. Densitometric quantification was performed manually using Fiji (gel analysis tool).

### Lentivirus production

For small scale production of lentiviral vectors containing shRNA constructs or the pLVX-expression plasmids, 3x10^5^ HEK 293T cells were seeded per 6-cm dish (2 ml media per well) 24 h before transfection. Transfection was performed using JetPEI (VWR International) with 1.5 μg of Vector DNA, 1 μg of psPAX2, and 0.5 μg of vesicular stomatitis virus G protein plasmid (pMD2.G) and 0.2 μg pAdvantage per well of a 6-well. Virus supernatants were harvested after 48 h, filtered through 0.45-μm-pore-size filters (Roth), and used immediately for transduction.

For the generation of stable T cell lines expressing the C5/C5L-mCherry constructs or primary human T cells expressing the Lifeact-GFP constructs, five 15 cm petri dishes were prepared with 2.5 x 106 HEK293T cells/dish in 22.5 ml medium. The transduction solution was prepared in a 50 ml reaction tube, containing: 112.5 μg vector, 40 μg (pMD2.G), 73 μg psPAX2, 25 ml NaCl and 500 μl JetPEI. The transduction solution was mixed and incubated at RT for 20min. For every dish 5ml of the solution was used. The dishes were incubated for 4 h at 37°C before changing the media. The supernatant containing virus particles was collected after 48h and filtered via 0.45 μm filter (Roth/Millipore). Virus was concentrated using 20% sucrose and ultracentrifugation at 24,000 rpm (Beckman SW28 rotor) for 2h at 4 °C. The supernatant was discarded and 200 μl fresh FCS free RPMI medium were added on the virus pellet and incubated for 30 min at 4 °C. The pellet was resuspended and stored at –80°C or directly used for transduction. Virus titers were assessed by determination of reverse transcriptase activity (SG-PERT).

### Transduction of human T cells

2-3x10^6^ JNLA or A3.01 cells were resuspended in 1.5E11 puRT/ μl concentrated virus solution or 1 ml of non-concentrated virus supernatant followed spin-transduction in 24-well plate format at 2300 rpm, for 1.5 h at 37°C, RT. After transduction the cells were incubated at 37 °C, overnight. The next day the cells were transferred into a 12-well plate and 3 ml complemented RPMI medium was added and incubated overnight. Cells expressing shRNAs or the C5/C5L-mCherry constructs were transferred to fresh medium 24h post transduction. 48h later puromycin (1.5μg/ml) was added and 72h post transduction, the medium was changed to fresh media with puromycin to accelerate cell growth. On day 4 post transduction, the cells were adjusted to the densities required according to the experimental question being addressed, with RPMI media without any selection antibiotics. Knock down was stable in the bulk culture for up to ∼1week post transduction. To generate stable A301 cells expressing either C5 or C5L tagged with N-terminal mCherry, the cells were FACS sorted for mCherry expressing cells post selection with puromycin for 1 week and then expanded in culture.

### Immunofluorescence Microscopy

As described previously (Tsopoulidis, Kaw et al. 2019), to study the actin dynamics in JNLA cells activated on stimulatory coverslips, 1-2x 10^5^ cells are put on the stimulatory coverslips for 5min at 37℃ before fixing them with 3% PFA. Following permeabilization and blocking, coverslips are incubated with primary antibodies overnight at 4℃ in 1% BSA(PBS). For phospho-specific targets/antibodies, all steps were done in 1X TBS. Following dilutions are used for the primary antibodies: rabbit anti-pTyr (1:100), rabbit anti-pSLP76 (1:1000) and mouse anti-mCherry (1:500). Species specific secondary antibodies conjugated to Alexa-Flour 568/647 (1:1000) were used for along with Phalloidin-Alexa Fluor 488 (1:600) for staining the F-actin. Although the nuclear lifeact reporter carries a nuclear export signal and thus also labels cytoplasmic F-actin, cortical F-actin is only labelled with low efficacy following permeabilization/fixation. An additional F-actin stain with Phalloidin is therefore required to efficiently stain and visualize cortical actin filaments. Coverslips were mounted with Mowiol (Merck Millipore) and analyzed by either epifluorescence microscopy (IX81 SIF-3 microscope and Xcellence Pro software; Olympus) or confocal microscopy (TCS SP8 microscope and LAS X software; from Leica).

### Generation of CRISPR-Cas9 based Knockout cells

We designed three single guideRNAs (sgRNA) for knocking out each of our gene of interest with the help of Synthego’s CRISPR design tool (https://design.synthego.com/#/). The sgRNAs were premixed with Cas9.3NLS (IDT) to create ribonucleoprotein complexes (RNPs) for faster and better editing efficiency as described earlier (Albanese, Ruhle et al. 2022). Premixed RNPs were then nucleofected (using either Amaxa 2b or Nucleofector 4D, Lonza) into the JNLA cells. JNLA cells post nucleofection are maintained in RPMI containing 10% FBS as a heterogenous pool, followed by knockout (KO) validation in the bulk pool using immunoblotting. For single KO clone expansion, nucleofected cell pools were expanded gradually for 4-5 days until they are validated for KO. Post KO validation, the cell suspension was diluted stepwise to reach a density of 0.5cells/50μl to seed 50μl/well in 96-U-bottom plates. Cells were then kept undisturbed for 1-2 weeks in the incubator until we observe change in color of the media. Wells with multiple colonies growing were discarded whereas single clonal populations were expanded gradually further until they were validated for KO using western blotting and surveyor assay, followed by initial functional characterization.

### Nuclear and cytoplasmic biochemical fractionation

JNLA cells were fractionated using the REAP method as described in (Suzuki, Bose et al. 2010). 8x10^6^ cells were harvested for each condition. The only difference we adapted is the manual sonication instead of an automated one. The number of sonication cycle varies between 10-15 (60s ON, 10s OFF) with the manual sonicator at 4°C or with ice. Each of the nuclear, cytoplasmic and total cell fractions are then immunoblotted as described above. As and when necessary, immunoblots were often stripped in 1X stripping buffer, followed by blocking for 1h at RT and re-probing with primary antibodies ON at 4℃. The protocol we followed for stripping including the preparation of stripping buffer were adapted from Abcam’s published protocol online (https://www.abcam.com/ps/pdf/protocols/stripping%20for%20reprobing.pdf)

## Supplemental information

### Supplementary Videos

**Suppl. Video 1: Live imaging of an IS formation between DMSO treated JNLA (in grey) and SEE treated B cells (in magenta).**

**Suppl. Video 2: Live imaging of an IS formation between CK869 treated JNLA (in grey) and SEE treated B cells (in magenta).**

**Suppl. Video 3: Live NFA and AR formation in DMSO treated Jurkat CD4 T cells upon falling on a stimulatory surface.**

**Suppl. Video 4: Live NFA and AR formation in CK869 treated Jurkat CD4 T cells upon falling on a stimulatory surface.**

### Supplementary Figures

**Suppl. Fig 1:**
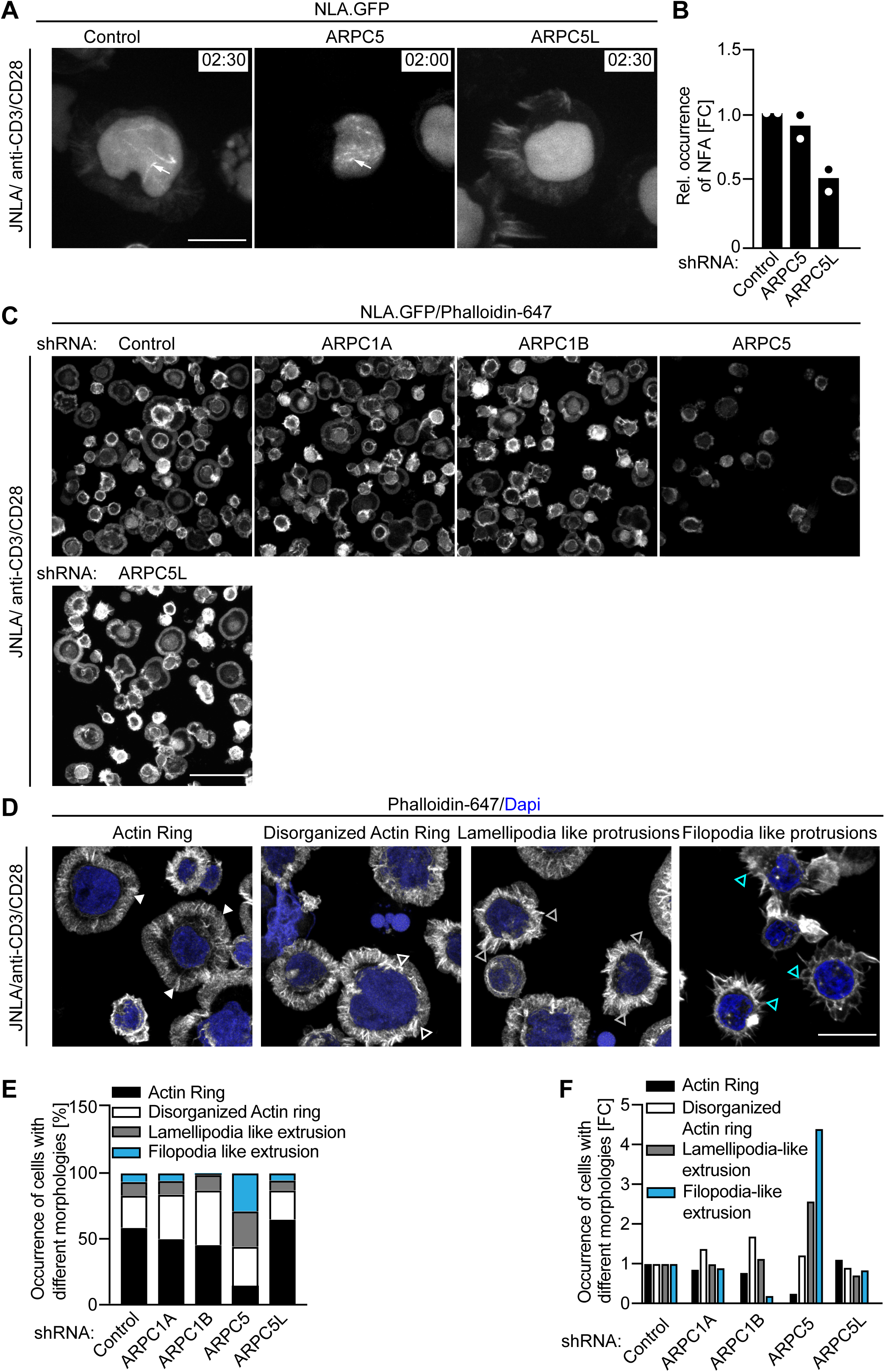
ARPC5 knockdown cells exhibit filopodia and lamellipodia like morphotypes upon TCR activation. **(A)** shRNA mediated knockdown of scrambled (control), ARPC5 or ARPC5L in JNLA cells were put on TCR stimulatory GBDs and subjected to live-cell microscopy. Shown are representative still images from the spinning-disk confocal microscope indicates the time of cells forming NFA after falling and contacting the GBDs. Movies were acquired every 30s for a total of 6 min. Arrows indicate the nuclear F-actin (NFA). (**B)** Quantification of nuclear nuclear actin filaments [NFA] were performed upon contact with TCR stimulatory surface. Data points indicate mean values from two independent experiments where 40-60 cells were analyzed per condition in each experiment. **(C)** Representative bigger field of view of average intensity projections of confocal images showing Phalloidin-647 stained F-actin ring (AR) formation in JNLA cells, treated with indicated shRNA upon activation on coverslips coated with antiCD3+CD28 antibodies. **(D)** Representative confocal still images (average intensity projection) of JNLA cells upon activation for 5min on stimulatory coverslip showing formation of distinct morphologies of actin ring formed. Cells were fixed, permeabilized and stained for F-actin (with Phalloidin 647) and counterstained with DAPI. **(E-F)** Cells with knockdown, exhibiting different morphologies (bars in different colours, stacked) based on classification shown above in (D), upon TCR activation, were quantified as % of cells of the total 100 cells quantified per condition is represented in (E). **(F)** shows the fold change in the different morphotypes exhibited by the cells upon knockdown relative to the control cells starts forming (top panel) or has fully formed into a classical actin ring structure (bottom). Scale bar, 7µm.

**Suppl. Fig 2:**
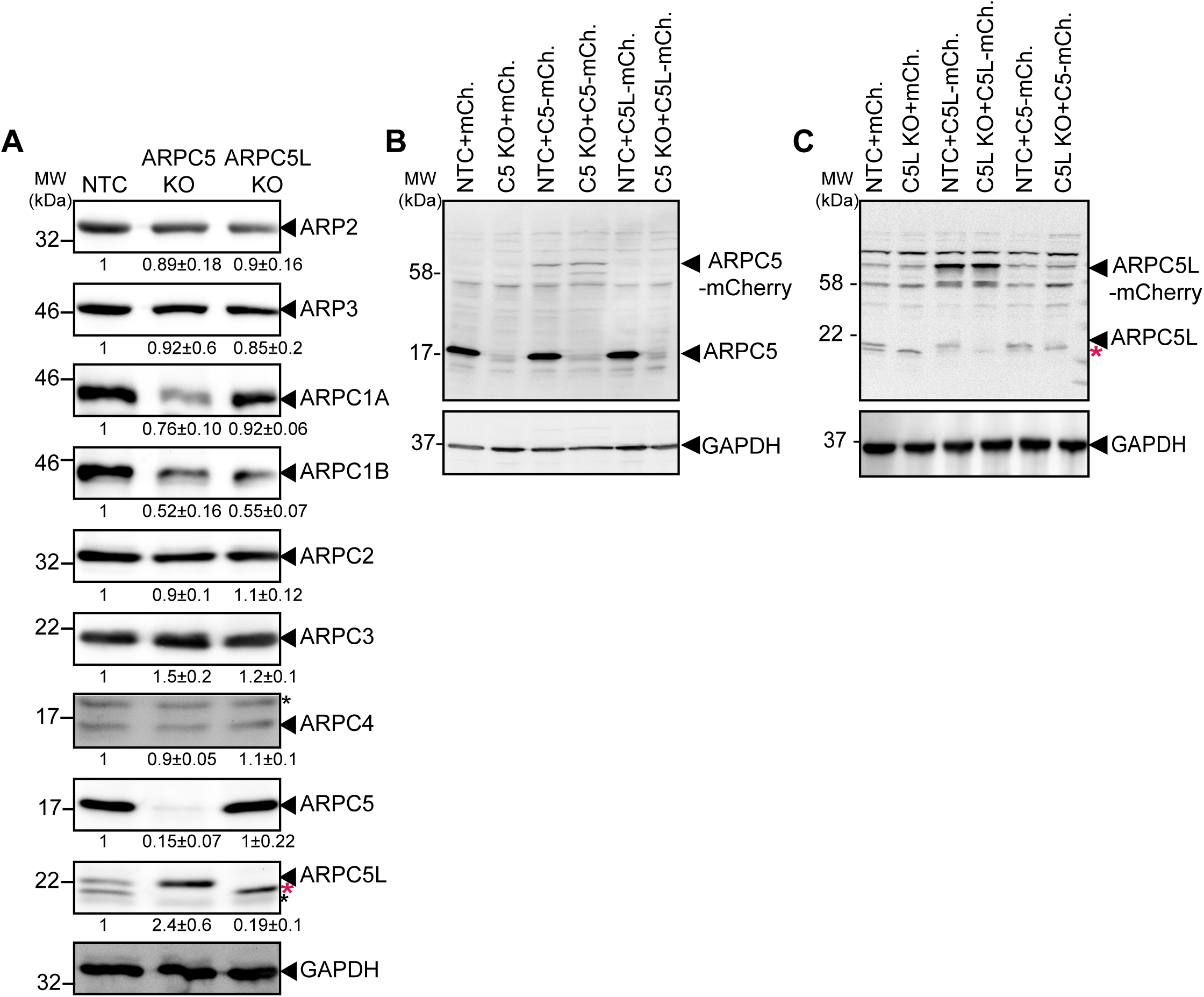
Expression of Arp2/3 subunits upon knockout of respective ARPC5 isoforms & validation of the overexpression of C5 isoforms in the bulk culture. **(A)** Representative immunoblots show levels of each of the Arp2/3 complex subunits in JNLA cells with the indicated knockout (KO) of each of the ARPC5 isoforms. Black arrowheads indicate the specific bands, black asterisks mark unspecific bands. Immunoblots are representative of three independent experiments. The numbers indicated below the respective immunoblots (only the blots where differences were observed are indicated) represent mean±S.D values from three independent experiments, based on the densitometric quantification of the bands, normalized to GAPDH & compared to the NTC protein levels (which is set to 1). **(B-C)** Shown are representative immunoblots confirming the successful overexpression of each of the ARPC5 isoforms in JNLA cells on the indicated knockout (KO) of each of the ARPC5 isoforms compared to the nontargeting control (NTC) cells. Note that the ARPC5L antibody also detects ARPC5 (marked by red asterisk).

**Suppl. Fig 3:**
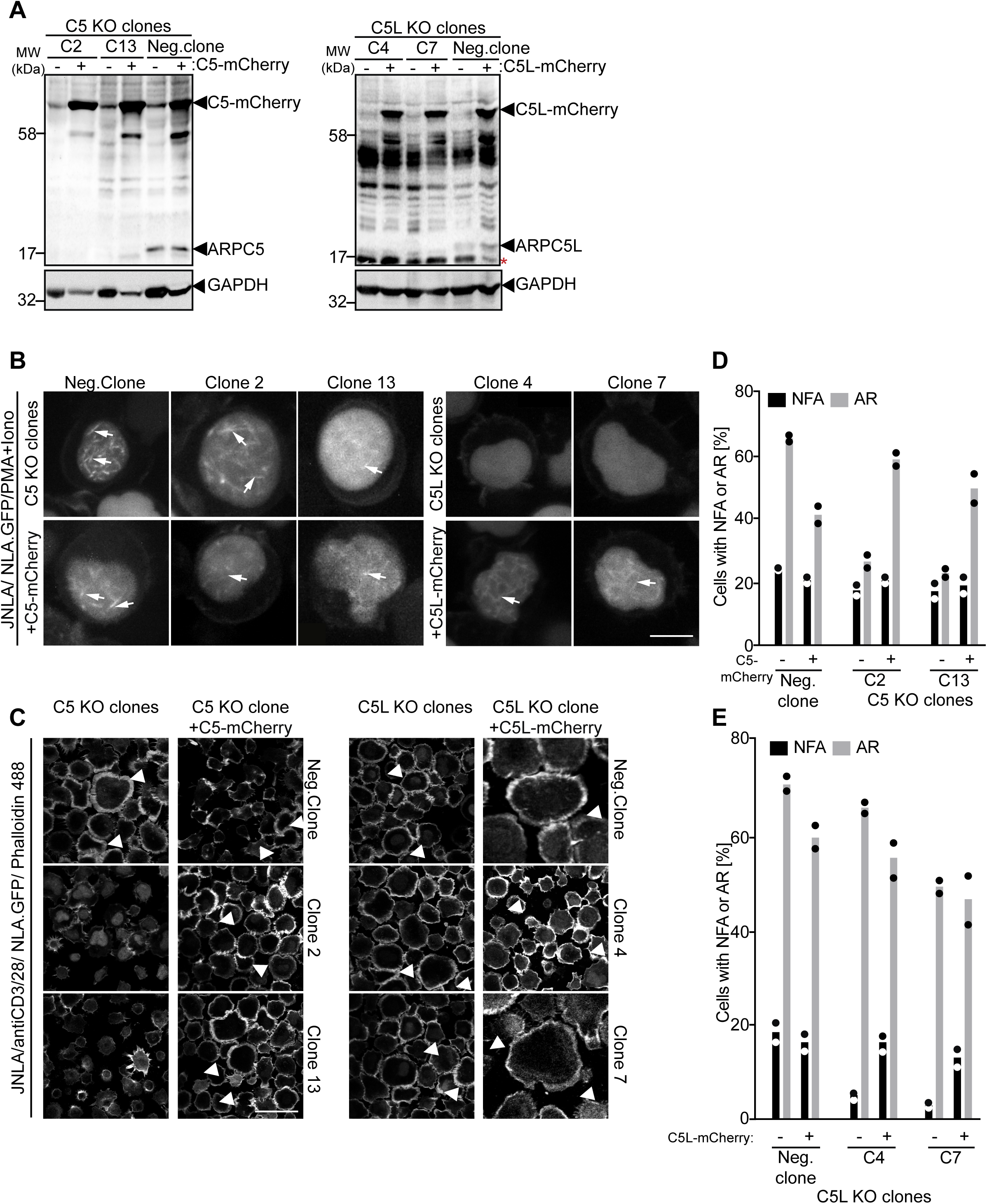
Rescue of the NFA and AR upon overexpression of the ARPC5 isoforms in JNLA knockout expanded from single KO clones. **(A)** Shown are representative immunoblots confirming the successful overexpression of each of the ARPC5 isoforms in JNLA cells on the indicated knockout (KO) clones of each of the ARPC5 isoforms compared to the clone where no knockout was observed (referred to as negative clone). **(B-C)** Shown are representative maximum intensity projections of confocal still images of the indicated KO clones overexpressing mCherry fusion proteins of the respective ARPC5 isoforms, post activation with either PMA+Ionomycin (P/I) as shown in (B) or on anti CD3+CD28 antibody coated stimulatory coverslips (as shown in (C). Arrows point to the nuclear F-actin (NFA) whereas arrowheads point to the F-actin ring. Scale bar, 7µm. **(D-E)** Quantification of NFA or AR formation in the respective KO clones or KO clones+ARPC5/C5L isoform expressing cells was performed indicating the % of cells forming NFA or AR. Bars indicate mean from two independent experiment where 30-40 cells were analysed for NFA and 100 cells were analysed for AR assay per condition in each experiment.

**Suppl. Fig 4:**
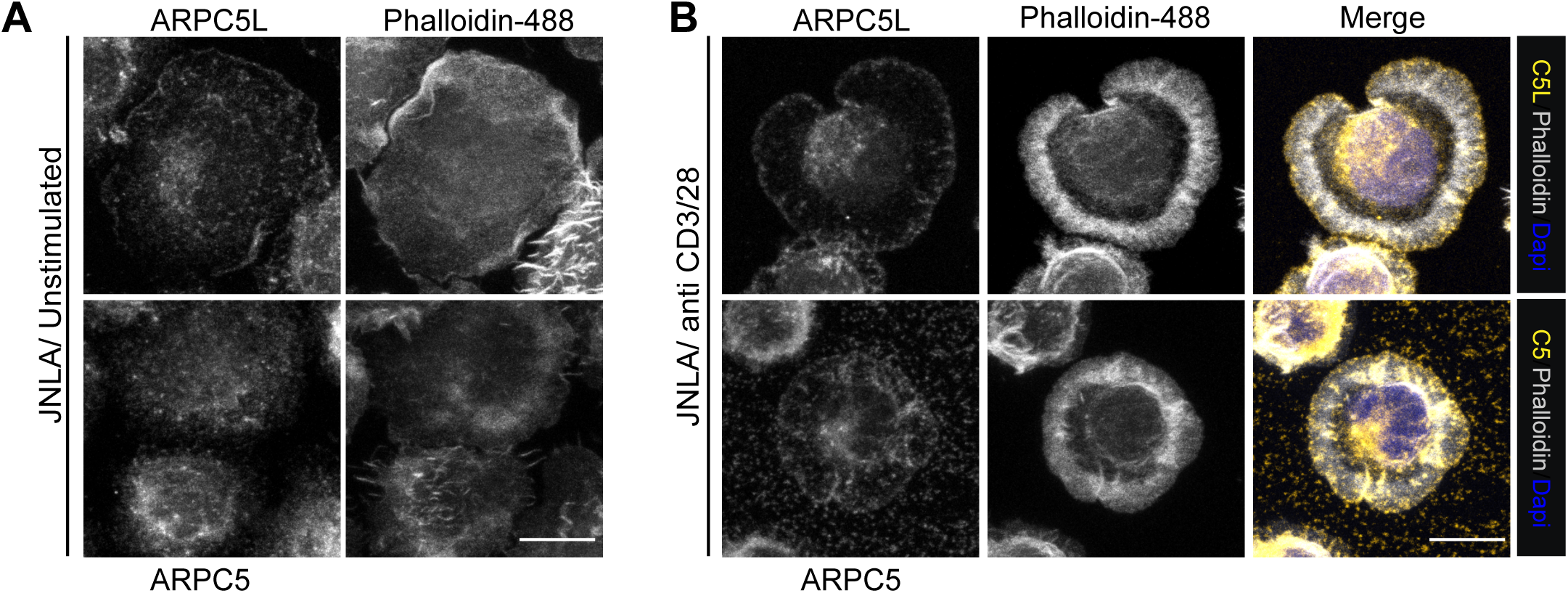
Subcellular distribution of ARPC5 isoforms. **(A)** Representative maximum intensity projection of confocal images showing respective ARPC5/ARPC5L antibody staining (on the left) of the endogenous ARPC5 isoforms expressed in the JNLA cells in unstimulated condition, where cells were fixed on poly-lysine coated coverslips. Additional, Phalloidin stained channel (on the right) is also shown for each image. **(B)** Representative maximum intensity projection of confocal images showing respective ARPC5/ARPC5L antibody staining of the endogenous ARPC5 isoforms, Phalloidin staining as well as merged view of DAPI, phalloidin and ARPC5/C5L antibody staining are shown in the JNLA cells upon TCR activation on coverslips coated with anti-CD3+CD28 antibodies. The high background observed in C5 antibody-stained channel is unavoidable as the CD3/28 antibodies coated on the coverslips and the ARPC5 antibody are of the same species (mouse). Scale bar, 10µm

**Suppl. Fig 5:**
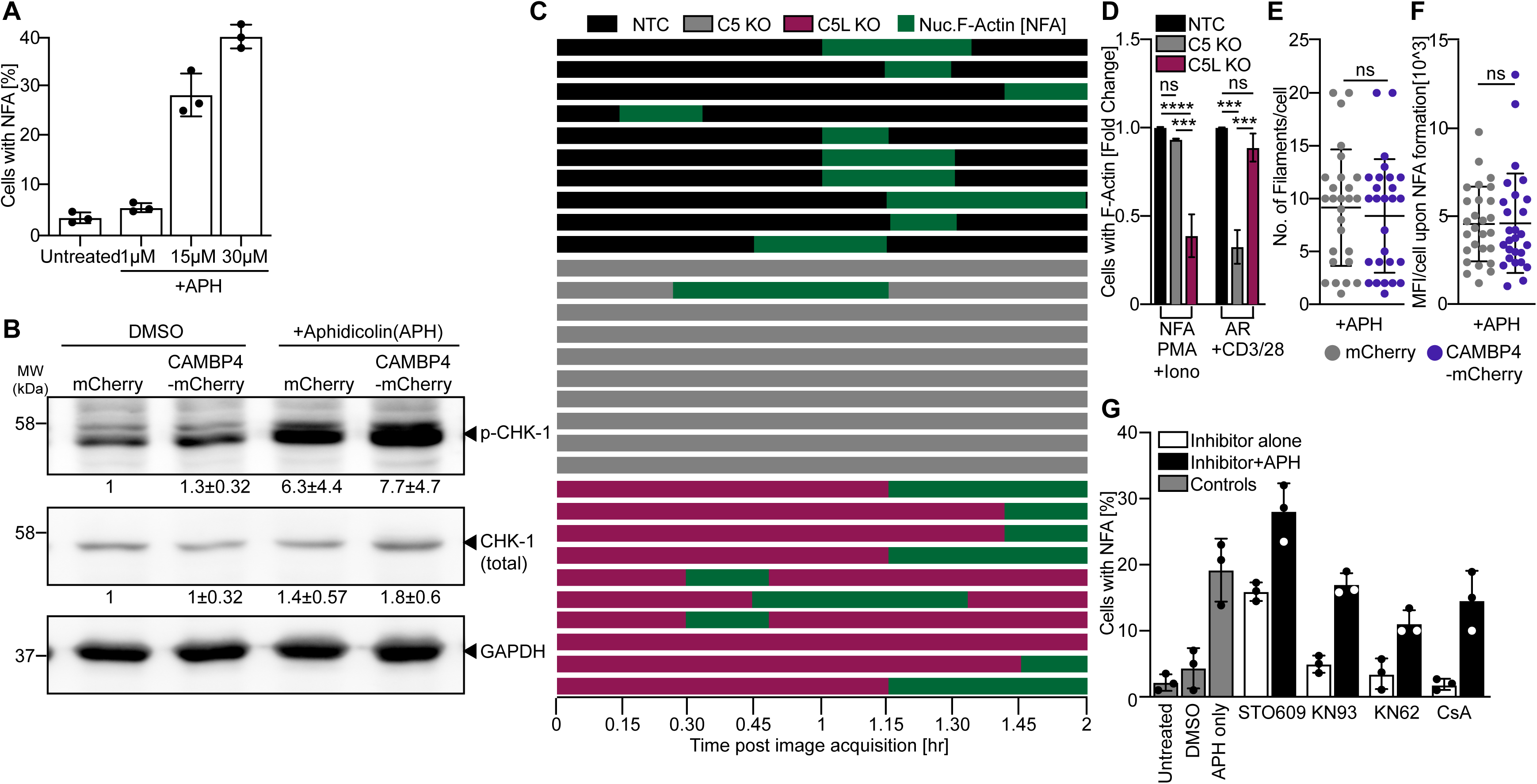
NFA formation induced by APH is not dependent on calcium signalling in CD4 T cells. **(A)** Bar graph represents mean of three experiments, showing % of cells forming NFA upon replication stress induction using different concentrations of Aphidicolin (APH). **(B)** Representative immunoblots shows induction of phospho-levels of DNA damage sensor CHK-1 in JNLA cells upon replication stress induction by Aphidicolin (15µM, APH) for 3h. Membranes were first probed with phospho specific antibodies, followed by stripping and re-probing with antibodies against total protein and GAPDH. Immunoblots are representative of three independent experiments where the numbers indicated below the blots represent mean±s.d intensity values of each condition when compared to mCherry as control (set to 1). Intensity values from densitometric analysis for both phospho & total protein levels were normalized to GAPDH before further comparisons were done. **(C)** Single cell tracking of 10 cells per condition (denoted by different colors for NTC, C5 or C5L KO) for the entire timeframe of 5h post pre-treatment with APH, shows the APH mediated NFA kinetics in KO and control JNLA cells. **(D)** shows relative comparison (Fold change, FC) of cells forming either NFA or AR, respectively, in Control and KO JNLA cells upon two different modes of T cell activation i.e., activation with PMA+Ionomycin (P/I) or on anti CD3/28 coated coverslips. Bars represent mean values from three independent experiments and error bars were calculated from mean±SD of 3 independent experiments where at least 30 cells were analyzed per condition per experiment for NFA quantification and more than 100 cells per condition per experiment were analyzed for AR quantification. Statistical significance was calculated using One-way ANOVA (Kruskal Wallis test) where *\*\*\*P ≤ 0.0002*, *\*\*\*\*P ≤ 0.000021* and ns: not significant. **(E-F)** Scatter plots represent the number of filaments per cell (E) and the intensity (MFI) per cell forming NFA (F) upon replication stress induction with APH comparing cells that express mCherry or nuclear specific CAMBP4-mCherry in JNLA cells. 20-30 single cells were analyzed manually using Fiji from each of the experimental conditions. **(G)** Blocking of the calcium signalling pathway downstream of Calmodulin, using inhibitors STO609, KN93, KN62 and Cyclosporin A (CsA) respectively on JNLA cells, followed by induction of replication stress with APH does not impair the replication stress mediated NFA burst observed. 30 cells/ condition were analysed in each experiment. Bar graph represents the mean from three independent experiments with each dot representing the mean per experiment.

**Suppl. Fig 6:**
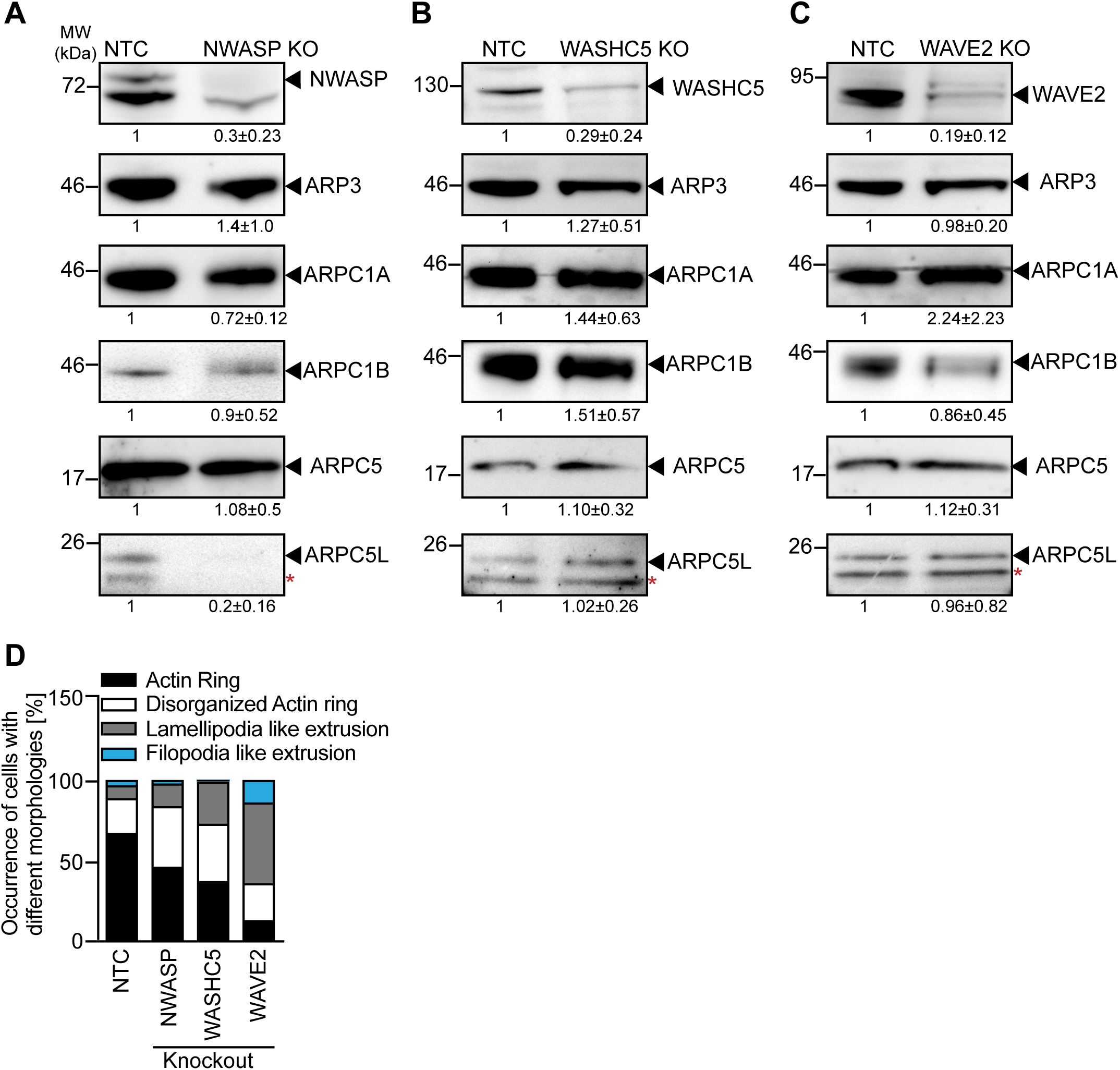
Molecular characterization of Arp2/3 complex upon NPF knockout in JNLA cells. **(A-C)** Representative immunoblots show levels of ARP3, ARPC1 & ARPC5 subunits in JNLA.GFP cells with the indicated knockout (KO) of respective class I NPFs. Black arrowheads indicate the specific bands, black asterisks mark unspecific bands. Note that the ARPC5L antibody also detects ARPC5 (marked by red asterisk). Immunoblots are representative of three independent experiments. The numbers indicated below the respective immunoblots represent mean±S.D values from three independent experiments, based on the densitometric quantification of the bands, normalized to Tubulin & compared to the NTC protein levels (which is set to 1). **(D)** Stacked bar graph represents the frequency of JNLA cells [%] with respective class I NPF knockout, exhibiting different morphologies based on classification shown in FigS2B, upon TCR activation, were quantified as % of cells of the total 100 cells quantified per condition.

